# Genetic and genomic architecture of species-specific cuticular hydrocarbon variation in parasitoid wasps

**DOI:** 10.1101/2021.10.18.464893

**Authors:** Jan Buellesbach, Henrietta Holze, Lukas Schrader, Jürgen Liebig, Thomas Schmitt, Juergen Gadau, Oliver Niehuis

## Abstract

Cuticular hydrocarbons (CHCs) serve two fundamental functions in insects: protection against desiccation and chemical signaling. CHC profiles can consist of dozens of different compounds and are considered a prime example for a complex trait. How the interaction of genes shapes CHC profiles, which are essential for insect survival, adaptation, and reproductive success, is still poorly understood. Here we investigate the genetic and genomic basis of CHC biosynthesis and variation in parasitoid wasps of the genus *Nasonia*. Taking advantage of the wasps’ haplo-diploid sex determination and cross-species fertility, we mapped 91 quantitative trait loci (QTL) explaining variation of a total of 43 CHCs in F_2_ hybrid males from interspecific crosses between three *Nasonia* species. To identify candidate genes, we localized orthologs of CHC biosynthesis-related genes in the *Nasonia* genomes. By doing so, we discovered multiple genomic regions where the location of QTL coincides with the location of CHC biosynthesis-related candidate genes. Most conspicuously, on a region on chromosome 1 close to the centromere, multiple CHC biosynthesis-related candidate genes co-localize with several QTL explaining variation in methyl-branched alkanes. The genetic underpinnings behind this compound class are not well understood so far, despite their high potential for encoding chemical information as well as their prevalence in both *Nasonia* CHC profiles and many other Hymenoptera. Our study considerably extends our knowledge on the so far little-known genetic and genomic architecture governing biosynthesis and variation of this fundamental compound class, establishing a model for methyl-branched alkane genetics in the Hymenoptera in general.

## INTRODUCTION

Understanding the genetic basis of quantitative phenotypic traits remains one of the central challenges in population genetics and evolutionary biology despite several advances in this field during the last decades (Yang et al. 2007; Mackay et al. 2009; Hartl 2020). Genes and regulatory nucleotide sequences governing natural variation in a set of phenotypic traits can be mapped (and potentially identified) in a genome by searching for phenotypic covariation with polymorphic genetic markers. This technique is commonly referred to as quantitative trait locus (QTL) mapping (Lander and Botstein 1989; Haley and Knott 1992; Mackay 2001a). Evolutionary genetic studies addressing variation and divergence of traits between closely related species with varying degrees of phylogenetic distance between them particularly benefit from QTL mapping techniques (Orr 2001; Slate 2005; Hartl 2020). Two components are of paramount importance for these kinds of studies: first, the availability of genetically tractable and crossable species with comparably short generation times and, ideally, established phylogenetic relationships (Mackay 2001b; Orr 2001; Mackay et al. 2009). Second, polygenic traits of interest that differ between the crossed species (Erickson et al. 2004; Slate 2005). Insects have proven to be particularly useful for QTL studies, since they combine various beneficial attributes: a multitude of quantifiable phenotypes, short generation times, large offspring numbers and, most commonly, relatively small genomes (Hoy 2005; Hunter and Kole 2008; Thomas et al. 2020). Cuticular hydrocarbons (CHCs), lipids constituting a major part of the waxy layer covering the insects’ epicuticle, are particular promising traits to dissect genetically via QTL mapping, as they are comparatively easily quantifiable, polygenic, and phenotypically as well as genotypically traceable (Chung and Carroll 2015; Blomquist and Ginzel 2021; Holze et al. 2021).

CHCs are long-chained molecules whose primary function is to protect terrestrial insects from desiccation (Ramsay 1935; Lockey 1980; Howard and Blomquist 1982, Fig. 1). As versatile semiochemicals, CHCs have also been demonstrated to function in a wide array of chemical communication systems integral to insect survival, reproduction, and adaptation (Carlson et al. 1971; Blomquist and Bagnères 2010; Blomquist and Ginzel 2021). Differences between CHC profiles arise from presence, absence, and abundance of individual CHCs, which can differ in their chain length as well as in the presence and position of double bonds and methyl groups (Lockey 1988; Howard and Blomquist 2005). The most commonly occurring compound classes are saturated straight-chain alkanes (*n*-alkanes), unsaturated straight-chain alkenes (*n*-alkenes) and alkadienes, and methyl-branched alkanes (Blomquist et al. 1987; Blomquist and Bagnères 2010). Despite considerable diversity of CHC profiles across insects, to the current stage of knowledge, the pathway of CHC biosynthesis appears to be mostly conserved (Blomquist and Ginzel 2021; Holze et al. 2021). Briefly, the biosynthetic pathway consists of the elongation of fatty-acyl-Coenzyme A units to produce very long-chain fatty acids (VLCFs) that are subsequently converted to hydrocarbons by subducting the carboxyl group (Nelson and Blomquist 1995; Howard and Blomquist 2005; Blomquist and Bagnères 2010, Fig. 1).

**Figure 1:**
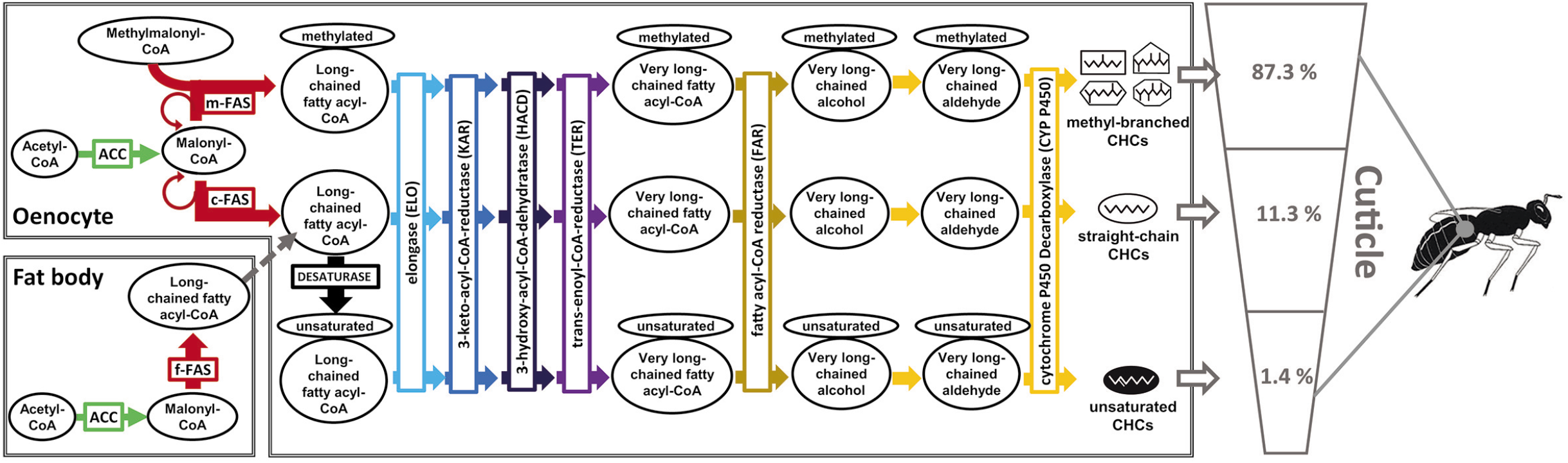
Simplified overview of CHC biosynthesis, with the main enzymes catalyzing the intermediate reactions highlighted (compare to Fig. 3-5 for the localization of the corresponding genes). The pathway branches at different stages, eventually synthesizing the different CHC compound classes that are then transported to the cuticle. The main CHC compound classes are methyl-branched alkanes (mono-, di-, tri-, and tetra-), straight-chain alkanes, and unsaturated alkanes. Acetyl-CoA as the initial reactant of CHC biosynthesis is mainly provided by the citric acid cycle. Respective percentages of the individual compound classes found on the cuticle of *Nasonia* males (averaged over our three study species *N. vitripennis*, *N. longicornis* and *N. giraulti*) are indicated as well. Schematic insect drawing depicting an *N. vitripennis* male adapted from Ward’s science, Rochester, NY, USA, 2002. Abbreviations: CoA: Coenzyme A, ACC: Acetyl-CoA carboxylase, FAS: Fatty acid synthase (m: microsomal, c: cytosolic, f: fat body associated). Adapted from Holze et al (2021).

Most of what we currently know about the genetics governing CHC biosynthesis and CHC profile diversity is based on studies on species of the Dipteran genus *Drosophila* (e.g. Jallon 1984; Coyne and Oyama 1995; Ferveur 2005). This taxonomic bias has so far impeded a more holistic view on the genetic underpinnings of this fundamental process across insect orders (Blomquist and Bagnères 2010; Holze et al. 2021). Particularly little is known about the genetic factors governing the differentiation and variation of methyl-branched alkanes, a dominant compound class in many insect taxa with high potential for encoding chemical information (Niehuis et al. 2011; Chung and Carroll 2015; Holze et al. 2021). In contrast to *Drosophila*, where methyl-branched alkanes only comprise a small fraction of their CHC profiles, this particular compound class dominates the CHC profiles in most species of the insect order Hymenoptera (Martin and Drijfhout 2009; Kather and Martin 2015; Sprenger and Menzel 2020). However, no single gene with a demonstrated impact on methyl-branched CHC biosynthesis and variation has been unambiguously identified in Hymenoptera so far. To address this knowledge gap, suitable model organisms are required to provide a basis for studying the genetic background of CHC production, variation, and divergence.

The parasitoid jewel wasp genus *Nasonia* (Hymenoptera: Pteromalidae) has emerged as a model system well-suited to study quantitative as well as polygenic phenotypic traits in the Hymenoptera. It combines ease of maintenance and handling, the possibility to generate hybrids between its four described species, haplo-diploid genetics, and a growing genetic tool kit (Gadau et al. 2008; Werren and Loehlin 2009; Werren et al. 2010). The *Nasonia* species complex consists of four species, *N. vitripennis*, *N. longicornis*, *N. giraulti*, and *N. oneida* (Darling and Werren 1990; Raychoudhury et al. 2010). Natural populations of *Nasonia* are all infected with different strains of intracellular *Wolbachia* bacteria, leading to bidirectional incompatibility in most interspecific crosses (Breeuwer and Werren 1990; Bordenstein and Werren 1998; Bordenstein et al. 2001). However, the *Wolbachia* infections can be cured (Richardson et al. 1987), rendering hybrids between all four species readily obtainable (Breeuwer and Werren 1990; Bordenstein and Werren 1998; Raychoudhury et al. 2010). This has proven to be particularly useful for QTL mapping of interspecific differences in a variety of phenotypic traits (Gadau et al. 2002; Gadau et al. 2008; Niehuis et al. 2008). An exemplary study comparing CHC variation through QTL mapping between two *Nasonia* species, *N. vitripennis* and *N. giraulti*, shed some first light on the genetic basis of *Nasonia* CHC profile differences (Niehuis et al. 2011). A study on CHC variation between all so far described *Nasonia* species established clearly distinguishable species- and sex-specific CHC blends (Buellesbach et al. 2013).

Here we dissect the genomic and genetic architecture of CHC variation in hybrid cross-comparisons between three *Nasonia* species. We present results from conducting QTL analyses on CHC variation in recombinant F_2_ hybrid males obtained by crossing *N. vitripennis*, *N. longicornis*, and *N. giraulti,* and from mapping CHC biosynthesis candidate genes. We additionally screened for selection signatures considering candidate gene orthologs from two more distantly related parasitoid wasp species within the family Pteromalidae: *Muscidifurax raptorellus* and *Trichomalopsis sarcophagae* (Fig. 2). Finally, we discuss how CHC profile divergence between the three *Nasonia* species can be explained phenotypically when considering the newly gained knowledge on the genetic architecture of CHC profile differences between them.

**Figure 2:**
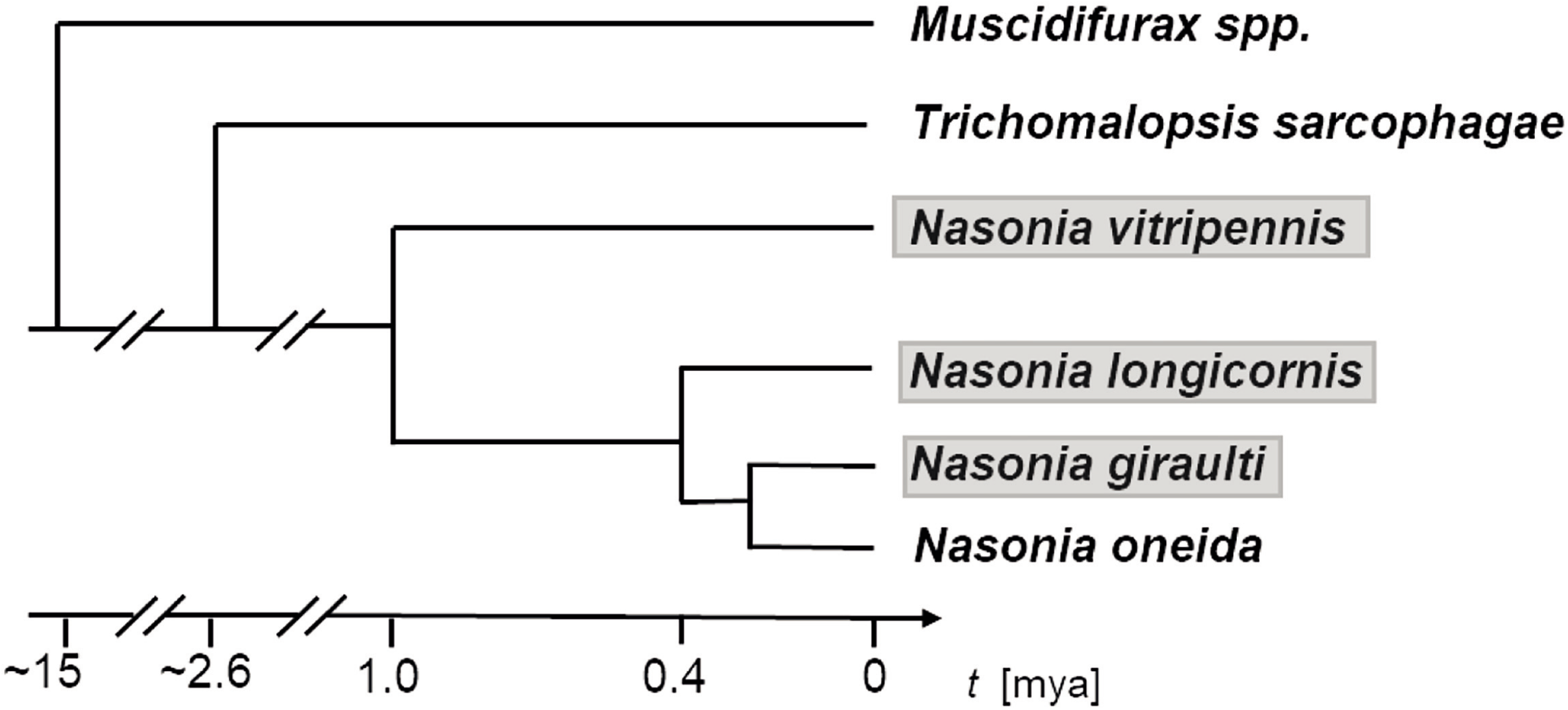
Phylogenetic relationship of the *Nasonia* species complex rooted by representatives of its two most closely related genera, *Trichomalopsis* and *Muscidifurax*, based on the D2 expansion region of their 28S rDNA sequences (Campbell et al. 2000) and mitochondrial DNA sequences (Werren et al. 2010). The three *Nasonia* species used in the current study are highlighted in grey.

## MATERIALS AND METHODS

### Nasonia strains and crossing experiments

We used the standard laboratory strains AsymCX, RV2x(U), and IV7(U) of the species *Nasonia vitripennis*, *N. giraulti*, and *N. longicornis*, respectively, to conduct the cross experiments (Breeuwer and Werren 1995; Werren et al. 2010). These strains are derived from wild-type strains and have been antibiotically cured from their *Wolbachia* infection, enabling them to hybridize under laboratory conditions. All strains were maintained in an incubator at 25 °C under permanent light and were provided pupae of the flesh fly *Sarcophaga bullata*. Cross experiments were conducted as described by Niehuis et al. (2008). We generated two types of F1 hybrid females: 1) by crossing virgin *N. vitripennis* females with *N. longicornis* males and 2) by crossing virgin *N. giraulti* females with *N. longicornis* males. All F1 hybrid females were kept unmated and thus generated recombinant (i.e., each genome constitutes a unique combinations of the parental genomes) haploid F_2_ males. These F_2_ recombinant hybrid males were collected 36 h after they had eclosed and were then freeze-killed and stored at −80 °C before characterizing their genotypes and CHC phenotypes.

### CHC profile analyses

We analyzed the CHC profiles of 100 *N. vitripennis* x *N. longicornis* (LV) and 101 *N. longicornis* x *N. giraulti* (LG) recombinant hybrid F_2_ males. We additionally characterized the CHC profiles of twelve *N. giraulti*, eleven *N. longicornis*, and 32 *N. vitripennis* males of the investigated strains for comparison. CHCs were extracted by submersing individual wasps for 10 min in 10 µl n-hexane (≥ 99.0%; Sigma-Aldrich, St. Louis, MO, USA) using 1-ml glass vials equipped with a 0.1-ml micro insert (Alltech, Deerfield, IL, USA). The CHC extracts were subsequently transferred to new vials and concentrated under a stream of nitrogen to approximately 1 μl total volume. The CHC extracts were then injected into a gas chromatograph coupled with a mass spectrometer (GC: 6890N; MS: 5975C; Agilent, Santa Clara, CA, USA) for analysis. The GC-MS was operated with in the splitless mode with an injector temperature of 250 °C. Separation of compounds was performed on a J&W DB-5MS fused silica capillary column (30 m × 0.25 mm ID; df = 0.25 μm; Agilent, Santa Clara, CA, USA) and applying the following temperature program: 60 °C start temperature, temperature increase by 40 °C per min to 200 °C, followed by an increase of 5 °C per min to 320 °C. Helium with a constant flow of 1 ml per min was used as carrier gas. The MS-quad temperature was 150 °C, the MS-source temperature was 230 °C, and the MSD-transfer line temperature was 315 °C. The obtained chromatograms were analyzed with the software Enhanced Chemstation E.01.00.237 (Agilent, Santa Clara, CA, USA). Peak identification was carried out automatically with the initial area reject parameter set to 0, the initial peak width set to 0.043, and the initial threshold set to 14. Shoulder detection was turned off. CHC compounds were identified according to their retention indices, diagnostic ions, and mass spectra (Carlson et al. 1998; Niehuis et al. 2011; Buellesbach et al. 2013).

### DNA extraction and QTL analysis

We extracted genomic DNA of all wasps after extraction of their CHCs. DNA extraction was done as outlined by Niehuis et al. (2007). Genotype data were collected for 29 length-polymorphic markers with known positions in the *Nasonia* nuclear genome (Tab. S1, Niehuis et al. 2011). The species-specific length of the markers was assessed by separating PCR-generated amplicons of them on a denaturing polyacrylamide gel following the procedure given by Niehuis et al. (2008). We used the following PCR temperature program for obtaining the amplicon: 5 min. at 95 °C, followed by 30 cycles of 1 min. at 95 °C, 1 min. at 55 °C, and 1 min. at 72 °C, and followed by 10 min. at 72 °C. PCR oligonucleotide primer sequences with their respective amplicon lengths are listed in Tab. S1.

Quantitative trait loci (QTL) analyses were conducted with the R/qtl package version 1.14-2 (Broman et al. 2003; Broman and Sen 2009) on a 64-bit-build of R version 2.10.1 (R Development Core Team 2018). Significant phenotypic differences in the relative abundances of CHC compounds were assessed with Bonferroni-corrected Wilcoxon rank-sum tests with continuity correction (Hollander and Wolfe 1999; Zar 1999). Each CHC compound was subjected to a one-dimensional one-QTL and to a two-dimensional two-QTL scan using Haley-Knott regression (Haley and Knott 1992) with an assumed genotypic error probability of 0.001 and a step width of 1 cM. We additionally applied a multiple QTL model to each CHC trait with forward/backward model selection as implemented in R/qtl. The maximum number of QTL for each trait was set to 10, with QTL positions refined after each step of the forward and backward selection. Penalties for model selection were set based on the permutation from the two-dimensional two-QTL genome scan of the respective CHC trait with a significance level of 0.01 (Manichaikul et al. 2009). Significance thresholds for QTL presence were estimated from 10,000 permutations of each respective phenotypic CHC abundance. Since our QTL mapping was based on the *Nasonia* linkage map inferred by (Niehuis et al. 2010) and on genome assembly 1.0 (Nvit 1.0, Werren et al. 2010), we transferred marker positions and QTL to the most recent high-resolution linkage map of *Nasonia* (Nvit 2.1, Desjardins et al. 2013) where we performed the mapping of candidate CHC biosynthesis genes (Tab. S1, see below).

Though the position of several markers had slightly shifted on the new linkage map, their respective initial linkage groups remained the same, enabling the re-assessment of the QTL positions by their relative position between the two closest markers. A fixed confidence interval of 20 cM was chosen for each QTL, since this distance was equivalent to the average distance between markers.

### Search for candidate CHC biosynthesis genes

Thirty-eight well-characterized candidate genes with a demonstrated impact on CHC variation via targeted knockdown studies were selected from *Drosophila melanogaster* (Tab. S2, see also Holze et al. 2021) via FlyBase, Version FB2019_02 (http://flybase.org/, Gramates et al. 2017). To screen for orthologs of these candidate genes in the *Nasonia* Nvit 2.0 reference genome, three complementary methods were used: first, the amino-acid sequences, including all possible isoforms encoded by *D. melanogaster* CHC biosynthesis genes of interest (GOI), were searched against the *N. vitripennis* proteome available on NCBI (*N. vitripennis*, https://www.ncbi.nlm.nih.gov/) using the BLAST+ software suite (version 2.8.1, Camacho et al. 2009). Amino-acid sequences with an e-value smaller than 10^-10^ were then again searched reciprocally against the *D. melanogaster* proteome. Reciprocal best hits were considered as orthologs. Second, the program OrthoFinder (version 2.3.1, Emms and Kelly 2015) was used to identify ortholog groups at the hierarchical level of the last common ancestor of the investigated species. OrthoFinder generated a) a list of ortholog groups that contained one gene from each of the tested species only (single-copy genes) and b) rooted phylogenetic trees of genes that belong to ortholog groups with paralogous gene copies. The software DIAMOND (Buchfink et al. 2014) was used to align the amino-acid sequences of a given ortholog group. Third, we queried the WaspAtlas database (http://cyverse.warwick.ac.uk:3000/, Davies and Tauber 2015) for ortholog groups (Flicek et al. 2014).

We were unable to identify unambiguous orthologs of some of the *D. melanogaster* CHC biosynthesis candidate genes in the *N. vitripennis* genome with the above-outlined methods despite partially high sequence similarities. Therefore, BLAST+-inferred hits in the *N. vitripennis* proteome with amino-acid sequence similarity thresholds of >-30-% identity, <-10^-10^ e-value, and >-85-% query coverage to *D. melanogaster* were additionally considered, although these may not necessarily constitute direct orthologs. The identification of orthologous candidate proteins of two protein families posed particular problems: fatty acid elongases (ELOs) and fatty-acyl-CoA reductases (FARs). Candidate ELOs and FARs in the *N. vitripennis* proteome with the highest amino-acid sequence similarity (lowest e-value and highest bit score) to the *D. melanogaster* ELOs and FARs with demonstrated impact on CHC variation (Tab. S2) were used as queries to search for homologs in the *Nasonia* proteome using blastp of the BLAST+ software suite. Hits above a similarity threshold of e-value < 10^-20^ were considered to belong to the same protein/gene family, and domain structure of the respective hits was taken into account to ensure that the derived gene sequences encode functional proteins. This was achieved with hmmscan of the HMMER software (http://hmmer.org/, version 3.1b2, February 2015) and use of the Pfam-A database (version 32, El-Gebali et al. 2019).

For visualization of the QTL regions and candidate gene positions on the *N. vitripennis* linkage map Nvit 2.1, the R package LinkageMapView was used (Ouellette et al. 2017).

### CHC profile discriminant analysis

Discriminant analysis (DA) was performed with the R package “MASS” (Venables and Ripley 2002) to test whether CHC profiles statistically differ between the six investigated groups of male wasps (i.e., those of the parental strains and those of the F_2_ hybrids). To standardize the peak area values for DA, the normalization method of the function “decostand” of the community ecology R package “vegan” was used based on the following formula (Dixon 2003):

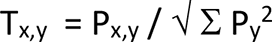

T_x,y_ is the transformed peak area x of individual y, P_x,y_ is the absolute peak area x of individual y and Σ Py^2^ to the squared sums of all absolute peak areas of individual y. This widely applied method for normalizing phenotypic data was chosen to make the peak areas comparable between our groups and to highlight the relative peak area differences. To visualize the data by plotting the first three discriminant functions simultaneously, the R package “scatterplot3d” was used (Ligges and Maechler 2003). Wilk’s λ was calculated to measure the quality of the DA.

### Gene orthology and screens for signatures of positive selection

To test for signatures of positive selection in our candidate genes, we extended our ortholog inference analysis to the fourth *Nasonia* species, *N. oneida*, as well as two closely related parasitoid wasp genera, *Trichomalopsis sarcophagae* and *Muscidifurax raptorellus*. The nucleotide sequences of the candidate genes from the *N. vitripennis* genome were extracted with SAMtools (Li et al. 2009) and aligned with the software BLAT (v35, Kent 2002; Bhagwat et al. 2012) to the nucleotide sequences of the latest genome assemblies of *N. giraulti*, *N. longicornis*, *N. oneida*, and *T. sarcophagae* kindly provided by Xiaozhu Wang (unpublished data). Aligned nucleotide sequences that shared at least 50 identical nucleotide positions and were aligned along at least 80 % query nucleotide sequence length were considered as homologous. To ensure these were also orthologous to the candidate genes of the *N. vitripennis* genome, their amino-acid sequences were reciprocally searched against the *N. vitripennis* proteome with BLAST+ (version 2.8.1, Camacho et al. 2009). Only genes that fulfilled the best reciprocal hit criterion were considered as orthologous. To annotate genes in the draft genomes of *N. giraulti*, *N. oneida*, *N. longicornis*, and *T. sarcophagae*, we used the program GeMoMa (version 1.6, Keilwagen et al. 2016). The reliability of the annotation by the program can be improved by providing it the coding sequence of an orthologous gene if a phylogenetically conserved gene structure can be assumed. We applied this approach for structurally annotating candidate genes in the *N. giraulti*, *N. oneida*, *N. longicornis*, and *T. sarcophagae* genomes, using nucleotide sequences of the orthologous *N. vitripennis* genes as references. Since BLAT was not sensitive enough to identify orthologs in *M. raptorellus* (comparatively distantly related to *N. vitripennis*, see Fig. 2), we *de novo*-inferred gene models in a draft genome of this species, kindly provided by Eva Jongepier (unpublished data), with the software GeMoMa. To obtain the respective coding nucleotide sequences from the GeMoMa-predicted annotations, the program BEDtools (Version 2.28.0, Quinlan and Hall 2010) with either the nucleotide sequences of the gene regions or the whole genome assembly.

Candidate genes were screened for signatures of positive selection by analyzing the ratio of nonsynonymous to synonymous substitutions (ω = dN/dS). The nucleotide sequences of each candidate gene ortholog group containing the orthologs of all six wasp species were aligned with the codon-sensitive alignment tool PRANK (version 170427, Löytynoja and Goldman 2008; Löytynoja 2014). The alignment process was supported with a phylogenetic tree (Fig. 2) of the study species whose evolutionary distances are based on the D2 expansion region of their 28S rDNA sequences (Campbell et al. 2000) and mitochondrial DNA sequences (Werren et al. 2010). Poorly aligned regions were trimmed with Gblocks (version 0.91b, Castresana 2000). Each sequence alignment was screened for signatures of positive selection on at least one site on at least one branch of the given phylogeny with the BUSTED (**B**ranch-**S**ite **U**nrestricted **S**tatistical **T**est for **E**pisodic **D**iversification) algorithm (Murrell et al. 2015), implemented in HyPhy (version 2.5, http://hyphy.org). Resulting p values were Benjamini-Hochberg-corrected for multiple testing (Benjamini and Hochberg 1995).

## RESULTS

### QTL explaining CHC variation in *Nasonia* F_2_ hybrid males

We detected a total of 91 QTL explaining CHC variation in recombinant F_2_ hybrid males. Of those, 80 were found in the recombinant F_2_ hybrid males obtained from crossing *N. longicornis* (♂) and *N. vitripennis* (♀) and eleven were found in the recombinant F_2_ hybrid males obtained from crossing *N. longicornis* (♂) and *N. giraulti* (♀) (Fig. 3 and 4, Tab. 1). The detected QTL explain variation in 43 (of a total of 53 analyzed) CHCs. The QTL of seven CHCs (two mono-methyl-branched, three di-methyl-branched, and two tetra-methyl-branched alkanes) were found in hybrids from both crosses, but corresponding QTL are located on different chromosomes or on different chromosomal regions (Tab. 1). The QTL of one single CHC compound, 15,17-DiMeC29 (RI: 2982), was detected exclusively in hybrids that originated from crossing *N. longicornis* and *N. giraulti* (Fig. 4 and Tab. 1). Sixty-nine QTL, explaining variation of 35 CHCs, were exclusively detected in hybrids that originated from crossing *N. longicornis* and *N. vitripennis* (Fig. 3 and Tab. 1). The QTL detected in F_2_ hybrid males of the latter cross are spread across all five chromosomes and encompass all six compound classes identified in *Nasonia* CHC profiles (i.e., alkanes, alkenes, mono-, di-, tri-, and tetra-methyl-branched alkanes). Several QTL explaining quantitative variation of structurally similar CHCs (mostly methyl-branched alkanes) clustered together, with two clusters each on chromosomes 1 and 2, and one cluster each on chromosomes 3, 4, and 5 (Fig. 3 and Table 1). Eleven QTL were found in recombinant F_2_ hybrid males obtained from crossing *N. longicornis* and *N. giraulti* (Fig. 2). All of them explain variation of methyl-branched alkanes (mono-, di-, and tetra-methy-alkanes). These QTL are located on chromosomes 2, 4, and 5, with clusters of them detected on chromosomes 4 and 5 (Fig. 4). Although these QTL clusters are in spatial proximity to the ones detected on the same two chromosomes in the F_2_ male hybrids obtained from crossing *N. longicornis* and *N. vitripennis*, they explain variation of different CHCs (compare Fig. 3 and 4).

**Figure 3:**
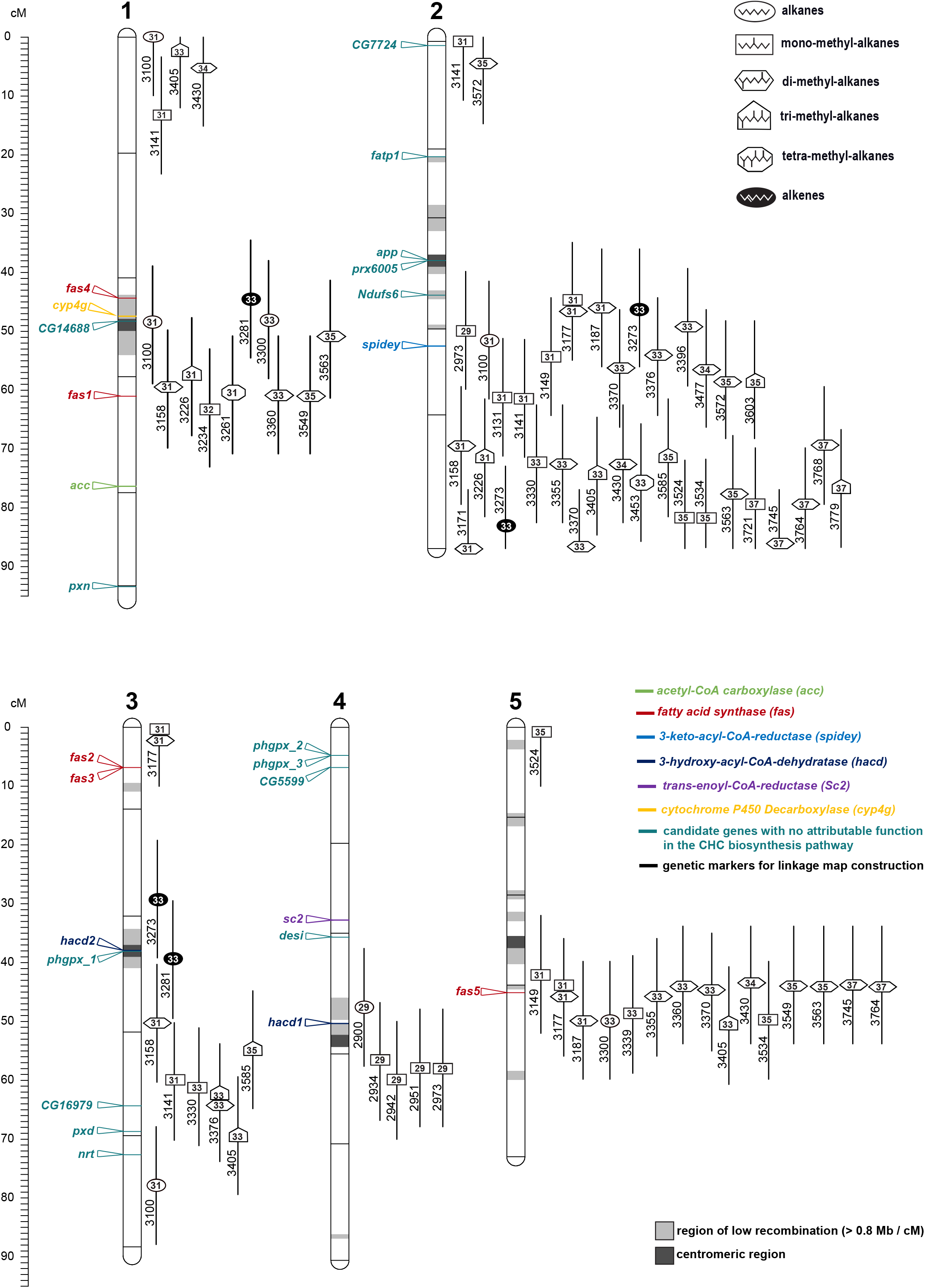
Linkage map based on F_2_ hybrid males from a cross between *Nasonia longicornis* (♂) and *Nasonia vitripennis* (♀) depicting positions of 80 quantitative trait loci (QTL) for 42 individual cuticular hydrocarbons (CHCs). Additionally, the positions of CHC biosynthesis-related candidate genes are shown. CHCs are indicated by symbols corresponding to their respective compound class (alkanes, alkenes, mono-, di-, tri-, and tetra-methyl-alkanes), their retention indices (RI) and numbers inside the symbols reflecting their carbon chain length. The five Nasonia chromosomes are labeled 1–5. Black horizontal lines on the chromosomes depict the respective positions of the 29 genotyped molecular markers used for determining the QTL positions. Orthologs to genes with a clear impact on CHC production derived from *Drosophila melanogaster* are color coded according to their putative position in the CHC biosynthesis pathway where possible (see Fig. 1). Regions of low recombination (> 0.8 Mb / cM) and centromeric regions are derived from Desjardins et al. (2013) and marked in light and dark grey, respectively.

**Figure 4:**
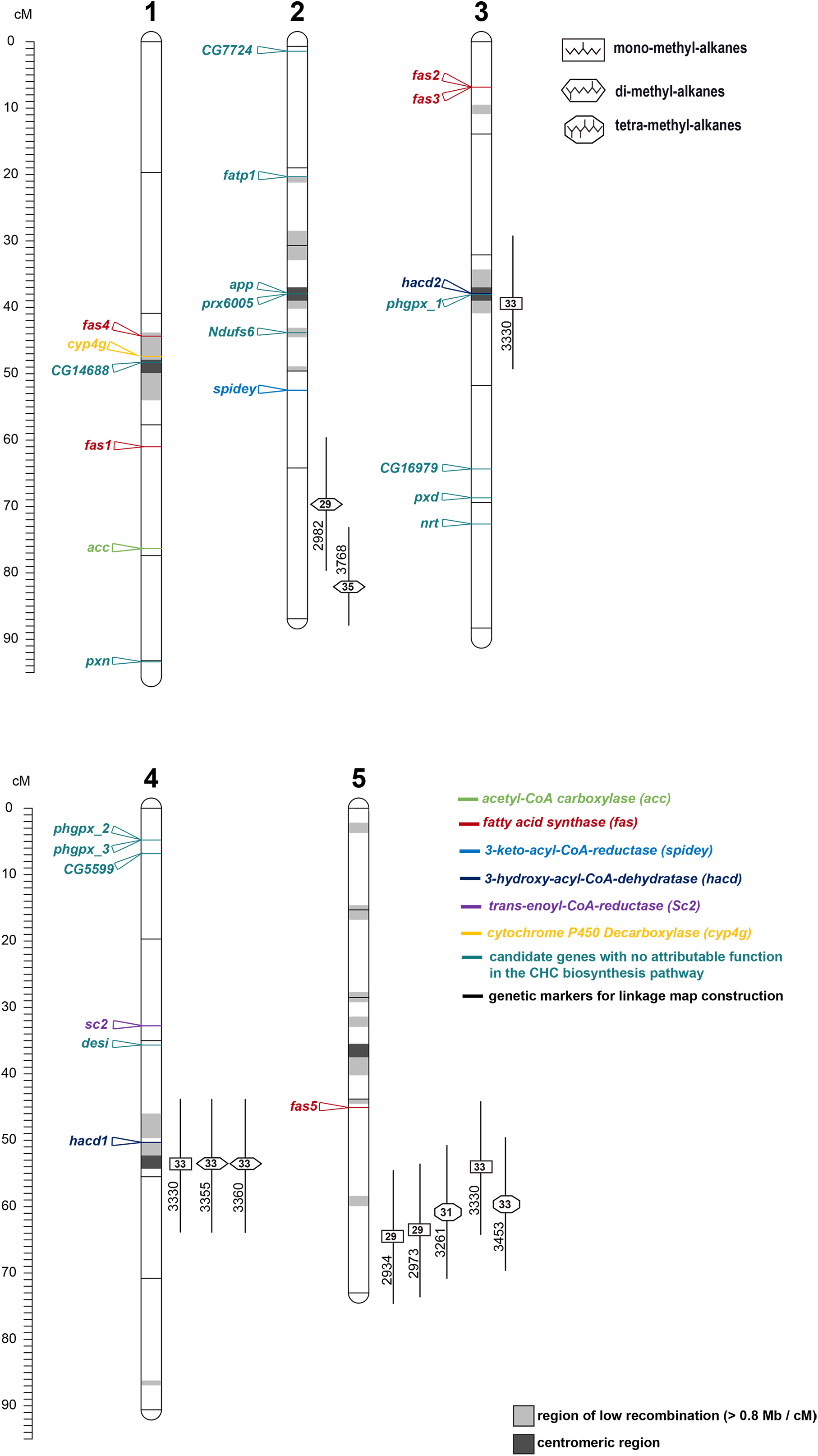
Linkage map based on F_2_ hybrid males from a cross between *Nasonia longicornis* (♂) and *Nasonia giraulti* (♀) depicting positions of 11 quantitative trait loci (QTL) for 9 individual cuticular hydrocarbons (CHCs). Additionally, the positions of CHC biosynthesis-related candidate genes are shown. QTL could only be detected for the variation of methyl-branched alkanes (mono-, di-, and tetra-methyl-alkanes). Labels of chromosomes, candidate genes, molecular markers and regions of low recombination are indicated as in Fig. 3

**Tab. 1:**
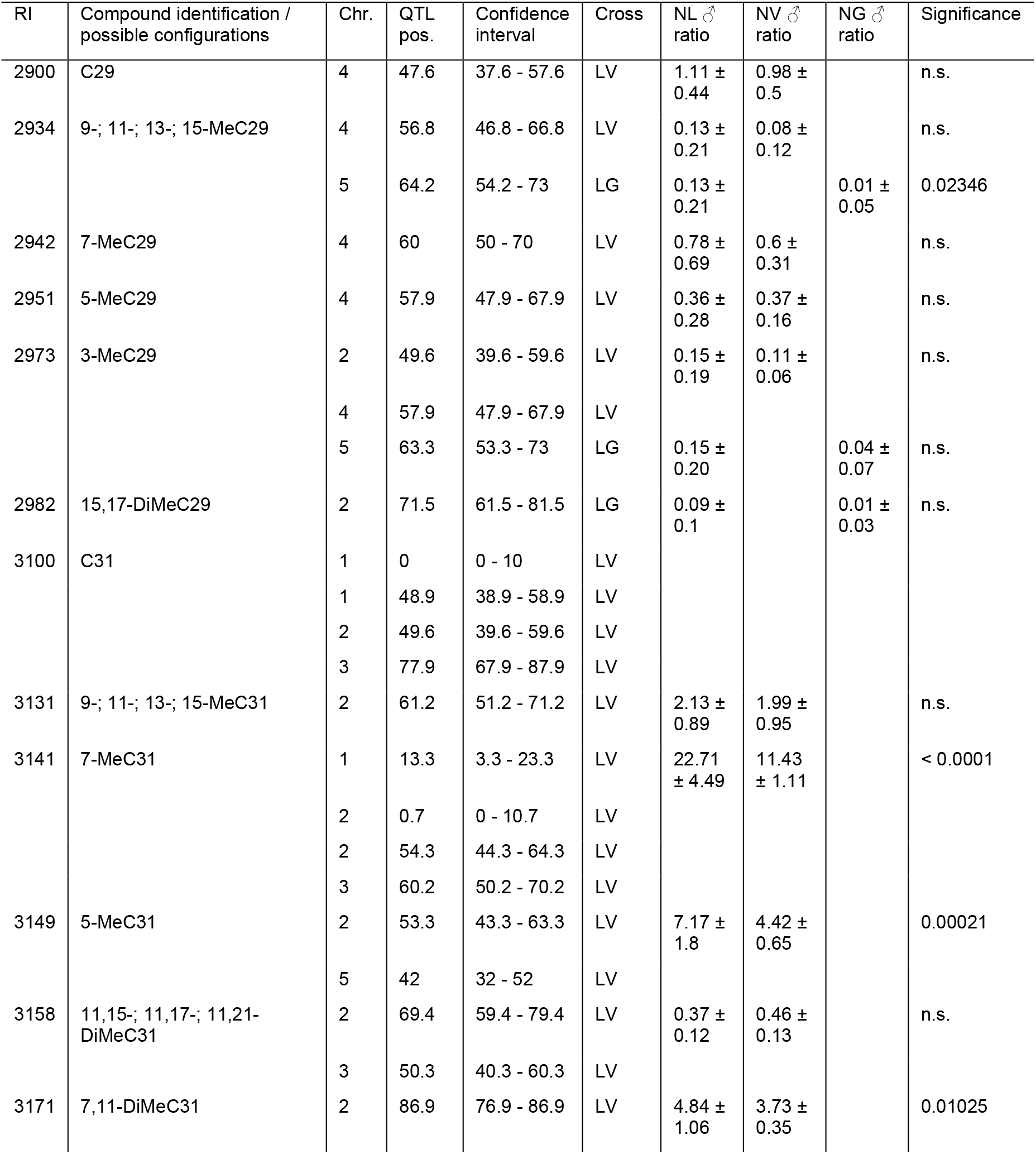
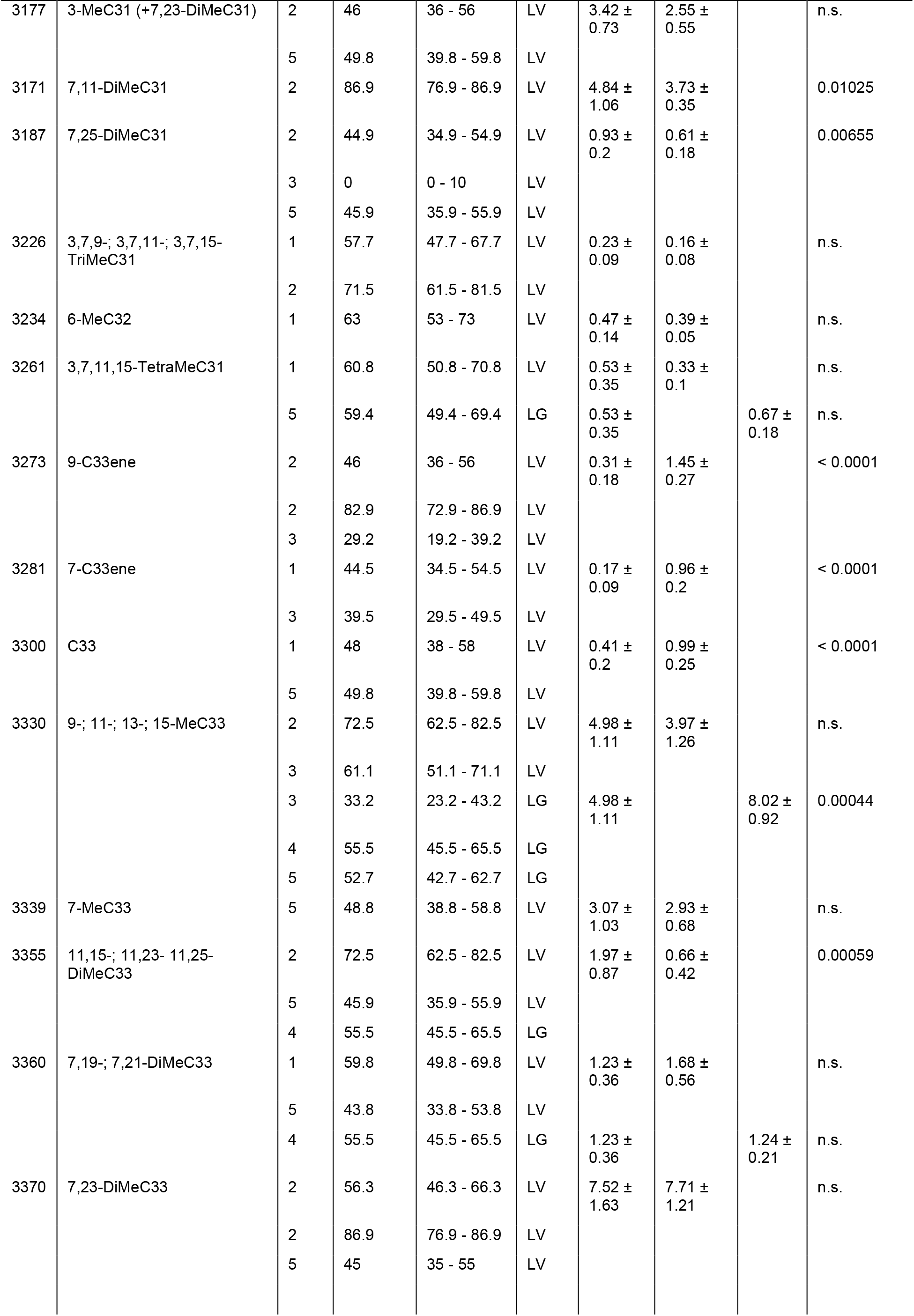
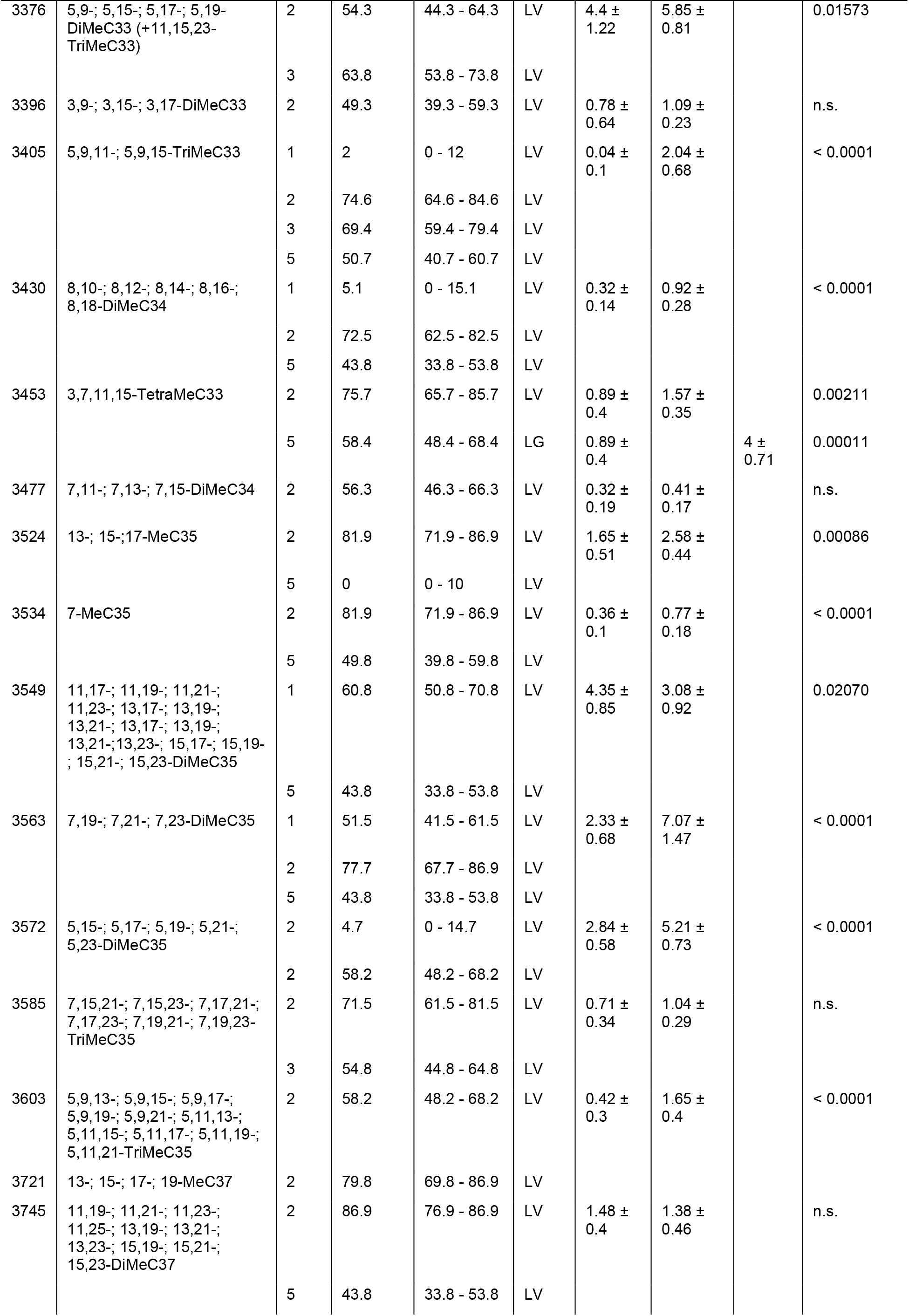
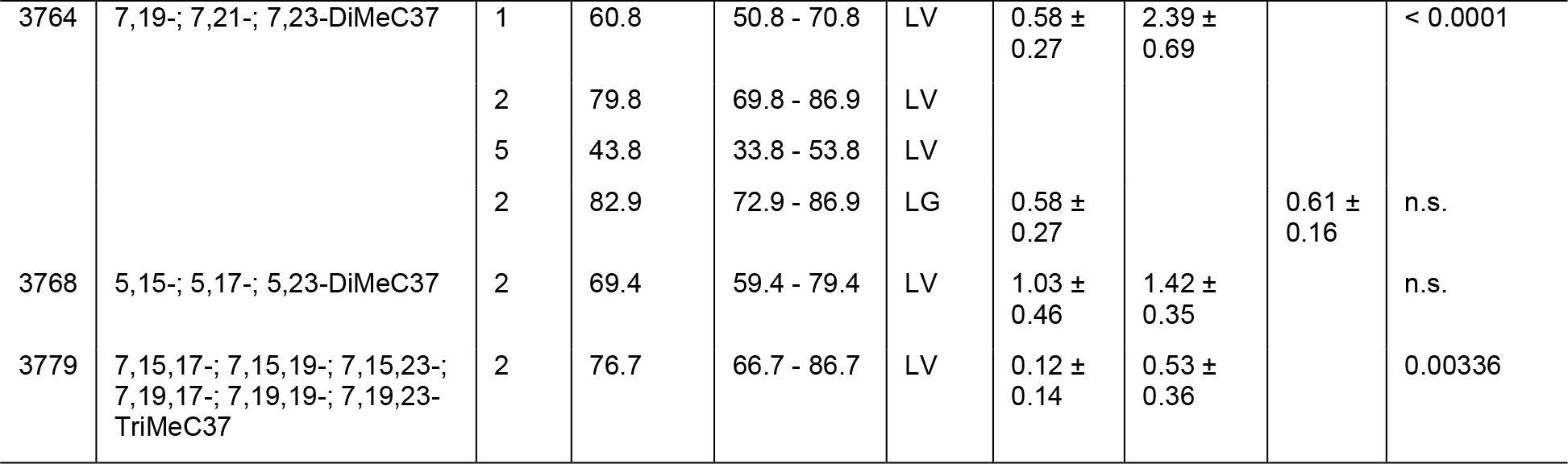
Cuticular hydrocarbons (CHCs) identified in male *Nasonia* profiles and the corresponding QTL from crosses between *Nasonia longicornis* and *Nasonia vitripennis* as well as between *N. longicornis* and *N. girauli*. Indicated are the retention indices (RI), compound identifications or possible configurations in case of ambiguities, chromosome number (Chr.), QTL position (pos.), confidence interval for the respective QTL position, hybrid cross (LV or LG), ratios of the CHC compounds in parental males of the two species for each respective cross (percentages with standard deviations) and significance assessments comparing the two ratios (Benjamini Hochberg-corrected Mann-Whitney U tests).

### CHC biosynthesis-related candidate genes

Of the candidate genes from *Drosophila melanogaster* with a demonstrated impact on CHC variation through targeted knockdown studies (Table S2), we found 15 to be represented by exactly one ortholog in the *Nasonia* reference genome, irrespective of the applied ortholog inference method (i.e., reciprocal BLAST search, OrthoFinder, and WaspAtlas) (Table 2).

**Tab. 2:** Candidate genes for CHC biosynthesis and variation in *Nasonia vitripennis* based on orthology towards *Drosophila melanogaster,* where the impact on CHC profiles of the respective genes has been demonstrated through targeted knockdown studies. Indicated are the gene annotations, gene IDs obtained from RefSeq, descriptions of the gene function obtained from NCBI, linkage map positions (centimorgan (cM) on the respective chromosome (c)), recombination rates (Mb/cM), and evidence of orthology based on the three methods reciprocal blast search (RB), OrthoFinder (OF) and WaspAtlas (WA). Genes for which exactly one ortholog could be unambiguously identified are indicated with a ‘+’, in other cases the method(s) that have generated the most consistently reliable results in each case are shown. Genes that can clearly be assigned to a particular step in CHC biosynthesis (compare to Fig. 1) are indicated in bold.

| Gene annotation | RefSeq ID | Description NCBI | Linkage map position | Recomb. rate (Mb/cM) | Evidence of orthology |
| --- | --- | --- | --- | --- | --- |
| **acc** | **100123347** | acetyl-CoA carboxylase | c1: 75.9 cM | 0.1114 | + |
| *app* | 100121627 | palmitoyltransferase ZDHHC18 | c2: 38 cM | 13.3806 | + |
| *CG14688* | 100120956 | phytanoyl-CoA dioxygenase | c1: 48.2 cM | 20.6198 | + |
| *CG16979* | 100119053 | ufm1-specific protease 2-like | c3: 64.2 cM | 0.1015 | + |
| *CG5599* | 100122501 | lipoamide acyltransferase | c4: 6.6 cM | 0.2967 | + |
|  |  | component of branched-chain alpha-keto acid dehydrogenase complex, mitochondrial |  |  |  |
| *CG7724* | 100120136 | 3 beta-hydroxysteroid dehydrogenase/Delta 5->4-isomerase type 4 | c2: 1.5 cM | 0.5931 | + |
| *Ndufs6* | 100119717 | NADH dehydrogenase (ubiquinone) Fe-S protein 6, 13kDa (NADH-coenzyme Q reductase) | c2: 43.8 cM | 1.1653 | + |
| ***cyp4g44*** | **100113546** | cytochrome P450 4G44 | c1: 47.5 cM | 2.6119 | WA |
| *desi* | 100115157 | uncharacterized LOC100115157 | c4: 35.8 cM | 0.1267 | + |
| ***fas1*** | **100121447** | fatty acid synthase | c1: 60.6 cM | 0.3184 | OF |
| ***fas2*** | **100122099** | fatty acid synthase-like | c3: 5.8 cM | 0.556 | OF |
| ***fas3*** | **100122083** | fatty acid synthase-like | c3: 5.8 cM | 0.556 | OF |
| ***fas4*** | **100119597** | fatty acid synthase-like | c1: 49.6 cM | 20.78 | OF |
| ***fas5*** | **100678737** | fatty acid synthase-like | c5: 45.3 cM | 0.7646 | OF |
| ***fatp1*** | **100120211** | long-chain fatty acid transport protein 4-like | c2: 20.4 cM | 0.8866 | RB |
| ***hacd1*** | **100119292** | very-long-chain (3R)-3-hydroxyacyl-CoA dehydratase hpo-8 | c4: 50.4 cM | 4.7456 | + |
| ***hacd2*** | **100115280** | very-long-chain (3R)-3-hydroxyacyl-CoA dehydratase | c3: 38 cM | 11.8001 | + |
| *nrt* | 100121165 | neurotactin | c3: 73 cM | 0.1823 | + |
| *phgpx_1* | 100677983 | probable phospholipid hydroperoxide glutathione peroxidase | c3: 38 cM | 11.8001 | RB,WA |
| *phgpx_2* | 100114603 | probable phospholipid hydroperoxide glutathione peroxidase | c4: 4.4 cM | 0.2203 | RB,WA |
| *phgpx_3* | 100123141 | phospholipid hydroperoxide glutathione peroxidase, mitochondrial-like | c4: 4.4 cM | 0.2203 | RB,WA |
| *prx6005* | 100117827 | peroxiredoxin-6 | c2: 38 cM | 13.3806 |  |
| *pxd* | 100119885 | peroxidase | c3: 68.6 cM | 0.3195 | OF,WA |
| *pxn* | 100116337 | peroxidase | c1: 93.4 cM | 0.1259 | + |
| ***sc2*** | **107980451** | very-long-chain enoyl-CoA reductase | c4: 32.1 cM | 0.1186 | + |
| ***spidey*** | **100121752** | very-long-chain 3-oxoacyl-CoA reductase | c2: 52.6 cM | 0.3081 | + |

Ortholog inference of the remaining 15 *D. melanogaster* candidate genes generated inconsistent results when comparing the three ortholog inference methods. For instance, reciprocal BLAST search (RBS) of three fatty acid synthase (FAS) genes with a demonstrated impact on *Drosophila* CHC biosynthesis identified nine putative *fas* orthologs in the *Nasonia* reference genome, whereas OrthoFinder found five. However, upon closer inspection, several of the putative *fas* orthologs identified by RBS are actually comprised of very short nucleotide sequences that unlikely encode fully functional proteins. These potential pseudogenes were excluded as CHC biosynthesis candidate genes in *Nasonia*. Furthermore, of all three ortholog inference methods, WaspAtlas was the only tool that led to the identification of an *N. vitripennis* ortholog to the *D. melanogaster* gene *cyp4g1*. However, due to a complimentary identification of an orthologous member of the *Cyp4g* gene family in the *N. vitripennis* genome with the highest nucleotide sequence similarity to *cyp4g1* by Qiu et al (2012), this ortholog was still included in the list of potential CHC biosynthesis candidate genes.

Two further gene families emerged in our analysis with high nucleotide sequence similarity to *Drosophila* CHC biosynthesis candidate genes despite no evidence for direct orthology: *elongases* (*elos*) and *fatty-acyl-CoA reductases* (*fars*). Of the four *elos* and the two *fars* in the *D. melanogaster* genome with a clear impact on CHC profile composition, we were able to identify twelve and 22 homologous genes with high nucleotide sequence similarity in the *Nasonia* reference genome (Tab. 3 and Fig. 5). Mapping of those genes on the *Nasonia* linkage map revealed a region on chromosome 1 around the centromere where eight *elos* and thirteen *fars* cluster together in close spatial vicinity (Fig. 5). Two additional candidate genes, orthologous to the *Drosophila* CHC biosynthesis genes *cyp4g1* and *CG14688*, also map in this chromosomal region.

**Figure 5:**
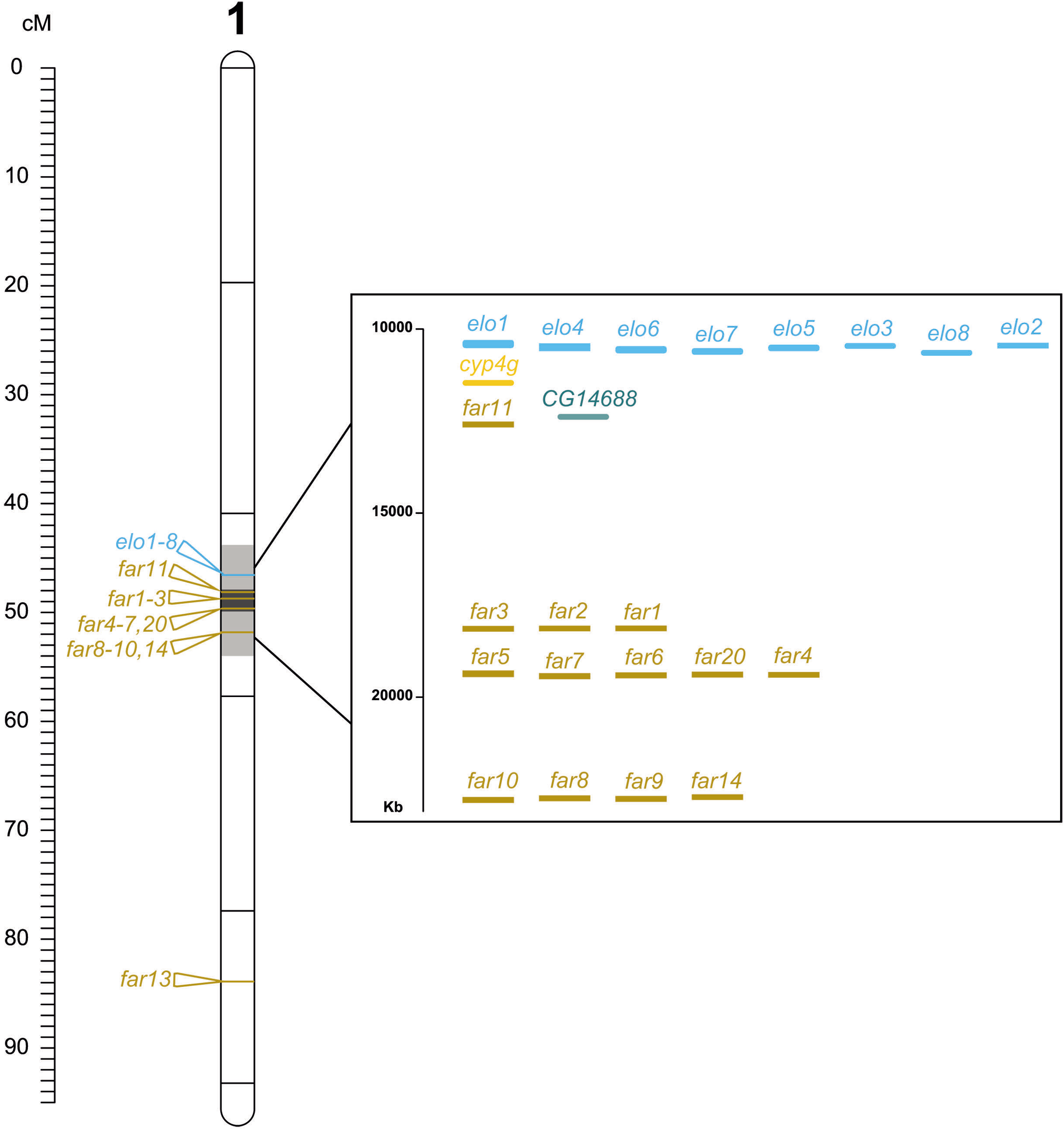
*Nasonia* linkage map of chromosome 1 with an enlarged region densely packed with CHC biosynthesis ted candidate genes. Eight elongase (elo) and thirteen fatty-acyl-reductase (far) cluster together in close spatial ity in addition to two additional genes genes cyp4g1 and CG14688 (see Tab. 2 and 3) in this chromosomal on. Note that this region maps around the centromere and is characterized by a particularly low recombination (> 0.8 Mb / cM). Labels of chromosomes and molecular markers are indicated as in Fig. 3 and 4.

**Tab. 3:** *Elongase* (*ELO*) and *fatty acyl-CoA reductase* (*FAR*) genes in the *Nasonia* reference genome homologous to *D. melanogaster ELOs* and *FARs* with a demonstrated impact on CHC profiles (*ELO*s: *CG18609*, *CG30008*, *CG9458*, *eloF*; *FAR*s: *CG10097*, *CG13091;* see Tab. S2). Indicated are the gene annotations, gene and respective protein IDs obtained from RefSeq, linkage map positions and recombination rates (Kb/cM).

| Gene annotation | RefSeq Gene ID | RefSeq Protein IDs | Linkage map position | Recombination rate (Kb/cM) |
| --- | --- | --- | --- | --- |
| *elo1* | 100114999 | XP_016845887.1<br>XP_008214189.1<br>XP_008214188.1<br>XP_001599838.1 | c1: 46.7 cM | 3.6211 |
| *elo2* | 100115035 | XP_001599867.1 | c1: 46.7 cM | 3.6211 |
| *elo3* | 100115068 | XP_003428006.1 | c1: 46.7 cM | 3.6211 |
| *elo4* | 100115103 | XP_008214193.1<br>XP_001599914.2<br>XP_016845633.1 | c1: 46.7 cM | 3.6211 |
| *elo5* | 100115131 | XP_001599942.1 | c1: 46.7 cM | 3.6211 |
| *elo6* | 100115199 | XP_001599996.2<br>XP_016838633.1 | c1: 46.7 cM | 3.6211 |
| *elo7* | 100115239 | XP_016838697.1<br>XP_001600017.1 | c1: 46.7 cM | 3.6211 |
| *elo8* | 100115278 | XP_001600048.1 | c1: 46.7 cM | 3.6211 |
| *elo9* | 100116202 | XP_008205141.1 | c3: 42.3 cM | 0.3664 |
| *elo10* | 100118619 | XP_008205429.1 | c4: 47.5 cM | 0.8598 |
| *elo11* | 100120094 | XP_001603768.2 | c3: 38 cM | 11.8001 |
| *elo12* | 100123468 | XP_001607111.2 | c5: 16.1 cM | 1.8931 |
| *far1* | 100115559 | XP_008213286.1 | c1: 48.9 cM | 22.4845 |
| *far2* | 100115595 | XP_008213287.2 | c1: 48.9 cM | 22.4845 |
| *far3* | 100115642 | XP_001600309.1 | c1: 48.9 cM | 22.4845 |
| *far4* | 100115711 | XP_001601168.1 | c1: 49.6 cM | 20.7846 |
| *far5* | 100117683 | XP_016845554.1<br>XP_016845553.1<br>XP_016845552.1<br>XP_016845551.1<br>XP_008205703.1 | c1: 49.6 cM | 20.7846 |
| *far6* | 100117801 | XP_008205709.1<br>XP_001601942.1 | c1: 49.6 cM | 20.7846 |
| *far7* | 100117837 | XP_001601970.1 | c1: 49.6 cM | 20.7846 |
| *far8* | 100118857 | XP_008212002.1<br>XP_008212003.1 | c1: 51.8 cM | 2.0759 |
| *far9* | 100118893 | XP_001602762.2 | c1: 51.8 cM | 2.0759 |
| *far10* | 100119003 | XP_001602857.1 | c1: 51.8 cM | 2.0759 |
| *far11* | 100121470 | XP_016841641.1<br>XP_016841638.1<br>XP_016841633.1 | c1: 48.2 cM | 20.6198 |
| *far12* | 100121766 | XP_008217013.1 | c5: 19 cM | 0.2367 |
| *far13* | 100123790 | XP_001607507.3 | c1: 83.2 cM | 0.5953 |
| *far14* | 103317077 | XP_016836513.1 | c1: 51.8 cM | 2.0759 |
| *far15* | 100117218 | XP_001601521.2 | c2: 85.4 cM | 0.1672 |
| *far16* | 100117254 | XP_001601550.4 | c2: 85.4 cM | 0.1672 |
| *far17* | 100117104 | XP_001601438.1 | c2: 85.4 cM | 0.1672 |
| *far18* | 100117143 | XP_001601466.1 | c2: 85.4 cM | 0.1672 |
| *far19* | 100117180 | XP_001601494.1 | c2: 85.4 cM | 0.1672 |
| *far20* | 100117760 | XP_016836801.1<br>XP_016836800.1 | c1: 49.6 cM | 20.7846 |
| *far21* | 100679544 | XP_003426741.2 | c2: 86.9 cM | NA |
| *far22* | 100117878 | XP_016843183.1 | c3: 38 cM | 0.1544 |

### Correlation in the genomic localization of candidate genes and QTL

QTL from the second cluster on chromosome 1 explaining variation of CHCs in recombinant F_2_ male hybrids of *N. longicornis* and *N. vitripennis* co-localize with CHC biosynthesis-related candidate genes in close spatial vicinity to the centromere (Fig. 3). Intriguingly, this also encompasses the conspicuous chromosomal region with a high density of *elo* and *far* genes (Fig. 5). Of the two detected QTL clusters on chromosome 2 from the cross mentioned above, the first one co-localizes with an ortholog of the candidate gene *spidey* that codes for a component of the elongase enzyme complex (Wicker-Thomas et al. 2015; Holze et al. 2021, see Fig. 1 and 3). Some QTL from the second cluster co-localize with several *far* genes (compare Fig. 3 and Fig. 5). On chromosome 3, the detected QTL cluster maps closely to the position of a gene orthologous to *CG16979*, a candidate gene with an impact on CHC variation in *D. melanogaster* despite an unclear role in the CHC biosynthesis pathway (Dembeck et al. 2015; Holze et al. 2021). On chromosome 4, several QTL map closely to a *hacd1*-ortholog, another gene that codes for an elongase enzyme complex component (Wicker-Thomas et al. 2015; Holze et al. 2021). On chromosome 5, a particularly large cluster, consisting almost entirely of QTL explaining variation in methyl-branched alkanes, co-localizes with a *fas* gene ortholog (*fas5*). Another QTL cluster identified in male F_2_ hybrids from the cross between *N. longicornis* and *N. giraulti* on chromosome 4 maps close to the *hacd1* ortholog, in close proximity to the above-mentioned QTL cluster from the cross between *N. vitripennis* and *N. longicornis* on the same chromosome (compare Fig. 3 and 4).

### CHC divergence between F_0_ parental and F_2_ hybrid males

To assess whether CHC profiles statistically differ between males of the parental strains and the recombinant F_2_ hybrid males, we conducted a discriminant analysis (DA) on the CHC profile data of 256 male wasps belonging to five groups: *N. giraulti* (NG)*, N. longicornis* (NL), *N. vitripennis* (NV), *N. longicornis* x *N. vitripennis* F_2_ hybrids (LV), and *N. longicornis* x *N. giraulti* F_2_ hybrids (LG) (Fig. 6). All groups differed in their CHC profiles significantly from each other (Wilk’s λ < 0.001, χ^2^ =23.22, p < 0.001), and the recombinant F_2_ males consistently clustered in between their respective F0 parental males.

**Figure 6:**
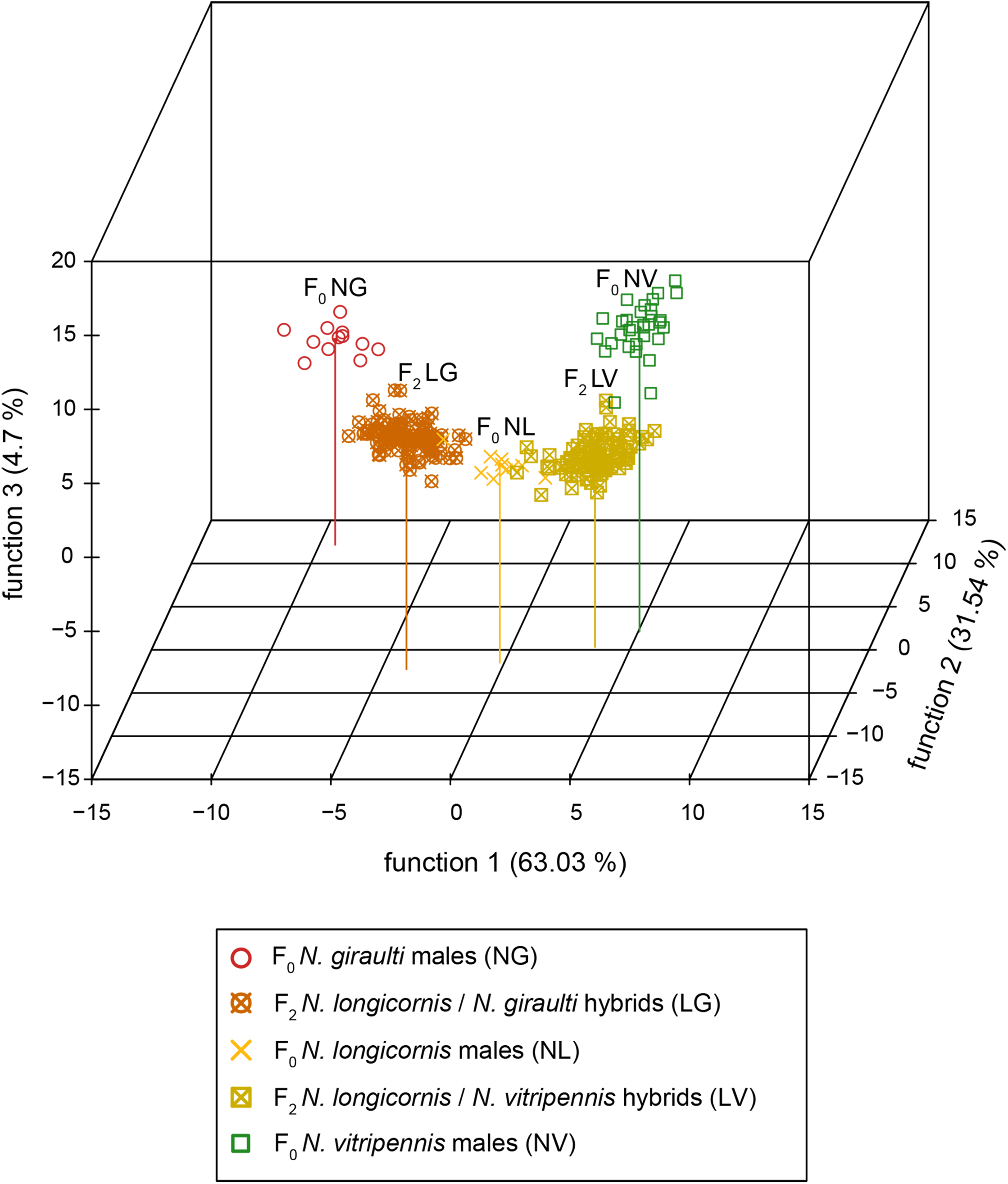
Plot of the first three discriminant functions showing the divergence between CHC profiles of F0 parental and F_2_ hybrid males between the three investigated *Nasonia* species in three dimensions. F0 NG: *N. giraulti* males (n = 12), F0 NL: *N. longicornis* males (n = 11), F0 NV: *N. vitripennis* males (n = 32), F_2_ LV: *N. longicornis* / *N. vitripennis* hybrid males (n = 101), F_2_ LG: *N. longicornis* / *N. giraulti* hybrid males (n = 100), total n = 256. All groups were significantly differentiated from each other (Wilk’s λ < 0.001, χ2 =23.22, p < 0.001), the variation each function explains is indicated in percentages.

### Orthology and signatures of positive selection for CHC candidate genes in other parasitoid wasps

We screened all 26 CHC biosynthesis-related candidate genes identified in the *N. vitripennis* reference genome (see Tab. 2) for orthologs in the respective genomes of the other three *Nasonia* species and in the genomes of two additional pteromalid wasp species: *Muscidifurax raptorellus* and *Trichomalopsis sarcophagae* (Fig. 2). In the genome of the latter, we were unable to identify orthologs of seven of the 26 *N. vitripennis* CHC biosynthesis candidate genes (*CG14688, fas2, fas4, fas5, hacd2, nrt*). Of these seven, two (*fas5* and *hacd2*) remained unidentified in *T. sarcophagae* and three remained unidentified in *N. giraulti* (*fatp1, nrt, hacd2*). Interestingly, *hacd2* was the only *N. vitripennis*-exclusive CHC biosynthesis-related candidate gene with no orthologs in any of the other investigated parasitoid wasp species (Tab. 4). Orthologs of the remaining 19 CHC biosynthesis-related candidate genes were found in all investigated species and were screened for signatures of positive selection based on the ratios of non-synonymous (dN) to synonymous (dS) nucleotide substitutions. After Benjamini-Hochberg correction for multiple testing, the signature for positive selection in one gene, *fatty acid synthase 1* (*fas1*), was considered statistically significant.

**Tab. 4.:**
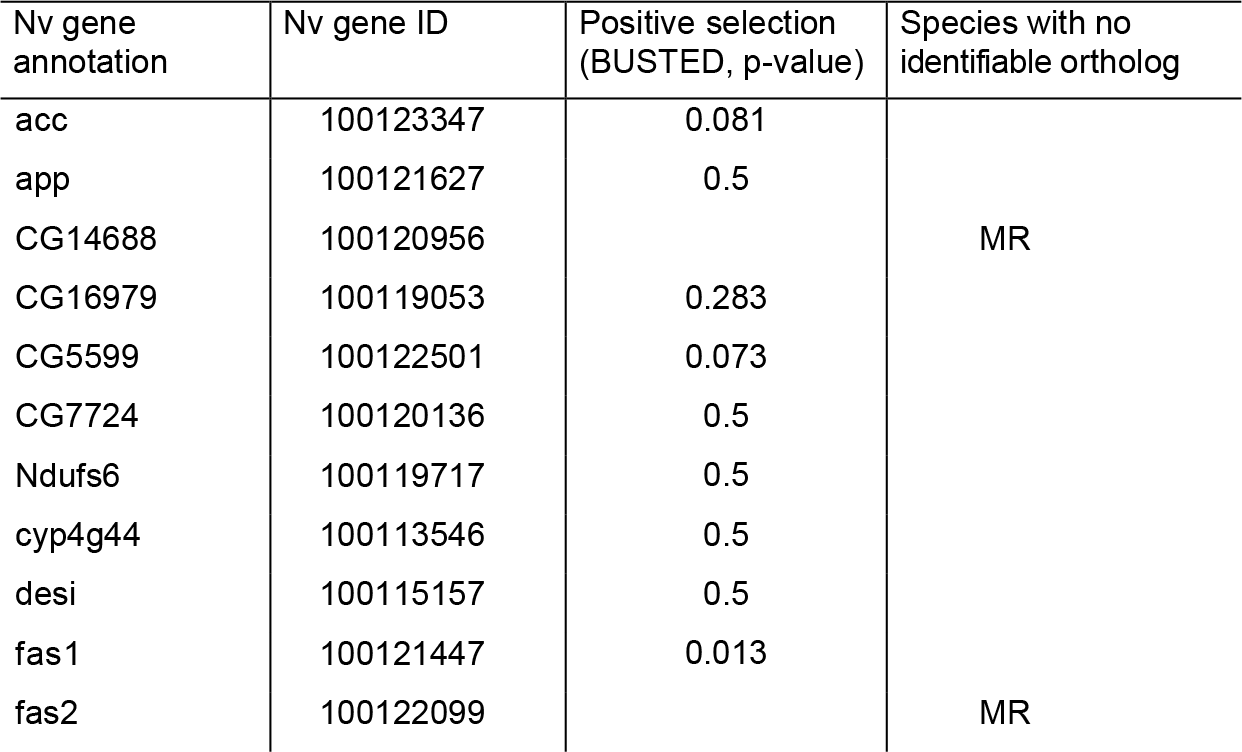
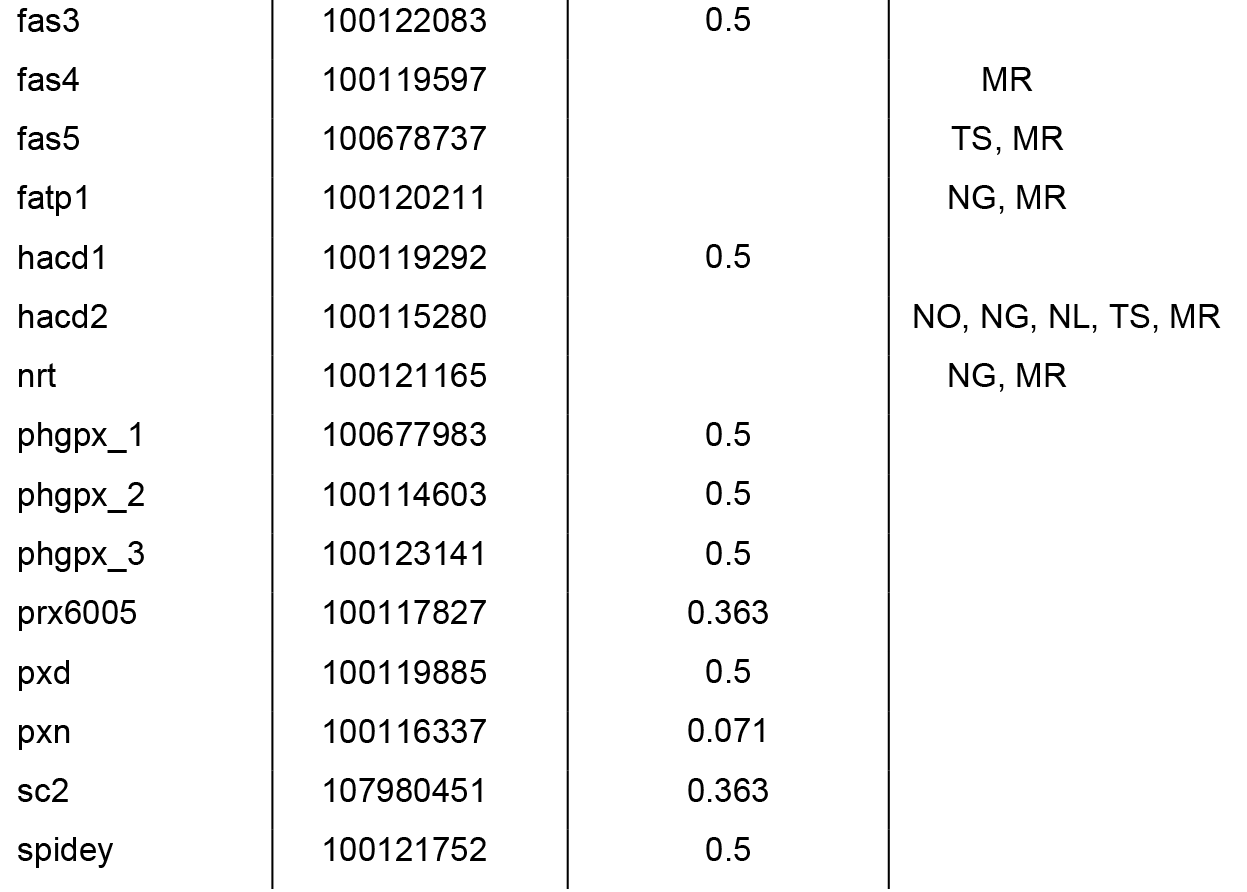
Orthologous candidate genes for CHC biosynthesis and variation in the *Nasonia* species complex as well as two representative species from the two most closely related genera, *Trichomalopsis sarcophagae* and *Muscidifurax raptorellus* (see Fig. 2 for phylogeny). Genes with identified orthologs in all analyzed species were subjected to a dN/dS ratio analysis to screen for signatures of positive selection with BUSTED. Indicated are gene annotations and gene IDs obtained from the *N. vitripennis* reference genome, species IDs where no orthologous gene could be unambiguously identified, and Benjamini-Hochberg-corrected p-values from the BUSTED analysis for signatures of positive selection. Species abbreviations: MR: *M. raptorellus*, TS: *T. sarcophagae*, NG: *N. giraulti*, NO: *N. oneida*, NL: *N. longicornis*.

## DISCUSSION

### Genomic regions harboring candidate genes and QTL explaining CHC variation

By analyzing recombinant F_2_ hybrid males from crosses between *N. longicornis* and *N. vitripennis* (LV) and between *N. longicornis* x *N. giraulti* (LG), we identified several QTL clusters mapping to genomic regions that harbor CHC biosynthesis candidate genes. Four of these regions additionally coincide with the location of QTL explaining CHC variation in recombinant F_2_ hybrid males of *N. vitripennis* and *N. giraulti* (VG, Niehuis et al. 2011, Fig. S1 and Tab. S3). The coinciding QTL of the latter study additionally explain variation in six CHCs (i.e., 7-MeC30, 5-MeC30, 9-C31ene, 3,7-DiMeC31, 3,25-DiMeC31, 3-MeC32, 15-; 17-; 19-MeC37) that we found no QTL of in the present investigation. Interestingly, QTL explaining variation of identical compounds in different hybrid crosses map only in very few instances to the same genomic regions (e.g., QTL for 11,15-; 11,23-11,25-DiMeC33 on Chr. 5 in LV and in VG crosses, QTL for 7,19-; 7,21-; 7,23-DiMeC37 on Chr. 2 in LV and in LG crosses). Conversely, in the majority of cases, QTL explaining variation of the same CHC compounds map to different chromosomal regions, or to different chromosomes entirely (Fig. 3, 4 and S1). This illustrates the genetic complexity and potential pleiotropy of the governance of most investigated CHC compounds. However, specifically comparing our cross between *N. vitripennis* and *N. longicornis* with the previously analyzed cross between *N. vitripennis* and *N. giraulti*, we identified overlapping QTL clusters governing the variation of structurally similar CHC compounds. The three most obvious of these clusters map close to the centromeric regions of chromosomes 1, 2, and 5 (compare Fig. 3 and S1, Tab. S3). The respective CHC compounds are all methyl-branched alkanes, partially sharing methyl-branching patterns and chain lengths (Tab. 1). Intriguingly, all of the above-mentioned QTL clusters co-localize with orthologs of candidate genes implicated with CHC biosynthesis. Particularly, the shared cluster on chromosome 1 mapped to the identified chromosomal region with the highest density of orthologs of CHC biosynthesis candidate genes (compare Fig. 3 and Fig. 5). As the recombination rate around the centromeres is very low, the close proximity of these gene orthologs is concordant with a higher likelihood of passing the whole unaltered gene cluster to the next generation, rendering gene shuffling in this region rare (Niehuis et al. 2010). The other two clusters co-localize with the positions of the gene orthologs *spidey* and *fas5* on chromosome 2 and 5, respectively. *Spidey* encodes a 3-hydroxy-acyl-CoA-dehydratase, constituting a part of the elongase enzyme complex (Wicker-Thomas et al. 2015), whereas *fas5* encodes a fatty acid synthase, instrumental for the formation of long-chain fatty acids as precursors for the elongation process (Juarez et al. 1992; Gu et al. 1997, see Fig. 2). These findings render these three distinct chromosomal regions particularly promising targets for further investigating the impact of the candidate genes mapped in these regions on CHC biosynthesis.

### CHC compound class variation and gene orthology between Nasonia and Drosophila

The vast majority of QTL that we detected in the hybrids from both crosses explained variation of methyl-branched alkanes (79 of 91 in total). This compound class dominates in CHC profiles of *Nasonia* wasp males and accounts for 87 % of the total amount of CHCs when averaged across all three investigated species (Fig. 1 and Tab. S4). This result is in accordance with those of most studies on CHC profiles in Hymenoptera, consistently reporting methyl-branched alkanes as the dominant CHC class (e.g. Martin and Drijfhout 2009; Kather and Martin 2015; Sprenger and Menzel 2020). In species of other insect orders, however, methyl-branched alkanes seem to be less dominant, which is particularly true for *Drosophila melanogaster*: analyzing CHC profiles in lines of the *D. melanogaster* Reference Panel (DGRP), Dembeck et al. (2015) detected only mono-methyl-branched alkanes in *D. melanogaster* CHC profiles, and these account for 16.2 % and 24.1 % of the total CHCs in CHC profiles of males and females, respectively (Tab. S5). This might be one of the reasons why our understanding of the genetics governing the variation in methyl-branched alkanes is comparatively limited.

The split between Hymenoptera and other holometabolous insects, including Diptera, is estimated to have occurred around 327 Mya (Misof et al. 2014). Thus, it is not surprising that were we unable to reliably identify orthologs of several genes with an impact on CHC profile composition in *Drosophila* in the genome of *Nasonia* wasps. For instance, the potentially best studied CHC biosynthesis-related genes in *Drosophila* are desaturases, catalyzing the introduction of double bounds in hydrocarbon chains (e.g. Wicker-Thomas et al. 1997; Coyne et al. 1999; Dallerac et al. 2000, Fig. 1). Although several *desaturase* genes have been identified in the *Nasonia* genome (Niehuis et al. 2011), we were not able to clearly confirm their orthology to *Drosophila* desaturases in the present study. However, the proportions of unsaturated CHCs vastly differ between the CHC profiles of *Drosophila* flies and those of *Nasonia* wasps (on average 59.2 % vs. 1.4 % in males, Tab. S4 and S5). Thus, since the proportion of unsaturated compounds is considerably less in *Nasonia* CHC profiles (compared to CHC profiles of *Drosophila* flies), the main genes governing their biosynthesis and variation might have deviated considerably after more than 300 Mya of estimated evolutionary divergence between *Nasonia* and *Drosophila* (Misof et al. 2014).

Similar in their unresolved orthology to *Drosophila*, we found numerous *elo* and *far* genes across the *Nasonia* genome (Tab. 3). In *D. melanogaster*, 19 *elo* genes have been identified in total (Chung and Carroll 2015; Zuo et al. 2018), but only four have been clearly associated with CHC biosynthesis and variation (Chertemps et al. 2007; Dembeck et al. 2015). Concerning *far* genes, from 17 identified in the *D. melanogster* genome (Finet et al. 2019), two have a demonstrated impact on CHC biosynthesis (Dembeck et al. 2015). This already indicates the difficulty in associating members in these two large and diverse gene families with CHC biosynthesis and variation, as they can also be involved in vastly different processes, including tracheal gas exchange, reproductive tissue growth, and larval development (Jaspers et al. 2014; Cinnamon et al. 2016). However, there are two further properties of *Nasonia* CHC profiles that greatly differentiate them from their *Drosophila* counterparts: first, in *Drosophila*, CHC chain lengths apparently do not exceed beyond C31 (Dembeck et al. 2015), whereas in *Nasonia*, CHC compounds with chain lengths of up to C37 have been identified, and additional CHCs with chain length of up to C52 have recently been described (Bien et al. 2019). Since elongases contribute to the elongation of long chain fatty acids (see Fig. 1), the longer and more diverse chain lengths in the *Nasonia* profiles could mean that a larger and more diverse set of *elongase* genes is functionally involved in CHC biosynthesis in *Nasonia* wasps. Second, *Nasonia* wasp CHC profiles appear to be more complex, generally containing more compound classes, including di-, tri-, tetra-methyl-branched alkanes, in contrast to CHC profiles of *D. melanogaster* (compare Tab. S4 and S5). It has been demonstrated that the *far* gene family shows particularly quick evolutionary turnover rates, which allows, in turn, for rapid diversification of CHC profiles between *Nasonia* species (Finet et al. 2019). This also argues for the involvement of a larger set of *far* genes in CHC biosynthesis in *Nasonia* than in *Drosophila* due to the more diverse and complex nature of *Nasonia* CHC profiles. Hence, the region on chromosome 1 containing eight *elo* and eleven *far* genes while harboring QTL mostly explaining methyl-branched alkane variation in two of our analyzed hybrid crosses constitutes a promising target-rich region for future functional genetic studies. *Nasonia* might prove to be a particularly well-suited model system for investigating the genetic underpinnings of methyl-branched alkane biosynthesis and diversification, with our current study providing a comprehensive foundation.

### Phylogenetic divergence in relation to CHC differentiation and selection signatures

It has been estimated that *N. vitripennis* has diverged around 1 Mya from the lineage from which the remaining *Nasonia* species evolved (Campbell et al. 1993). The divergences of the remaining *Nasonia* lineages occurred approximately between 500 000 and 400 000 years ago (Werren et al. 2010, Fig. 2). We discovered the smallest number of QTL governing CHC variation between the more recently diverged species pair studied by us: *N. longicornis* and *N. giraulti*. This finding matches well with the considerably shorter evolutionary time frame to accumulate genetic differences compared to the longer divergence time between each of the two species and *N. vitripennis*. Therefore, it is all the more surprising that phenotypically, the overall CHC profile divergence sufficiently discriminates all species with F_2_ hybrid males clustering as intermediate phenotypes between the respective F0 males (Fig. 6). As *N. longicornis* and *N. giraulti* males also differ considerable in their CHC composition from each other, this obvious phenotypic divergence is not reflected in the comparably low genomic divergence between these two species hinted at by our QTL comparison. A potential explanation for this might be other mechanisms, *e.g.*, phenotypic plasticity (Otte et al. 2018) or epigenetic regulation (Morandin et al. 2019), that factor into CHC profile divergence between these phylogenetically less distant species.

Concerning CHC biosynthesis candidate gene orthology, we extended our analysis beyond the genus *Nasonia* to two more distantly related parasitoid wasp species in the family Pteromalidae, *Trichomalopsis sarcophagae and Muscidifurax raptorellus* (Fig. 2). Unsurprisingly, in the genome of the latter, most distantly related species, we detected the fewest orthologs in comparison to the *N. vitripennis* reference genome (19 out of 26, Tab. 4). Furthermore, taking into account all shared 19 candidate genes in our six investigated wasp genomes, we detected signatures of positive selection in only one of them, *fas1*. The gene *fas1*, an ortholog to *FASN1* and *FASN2* in *Drosophila*, plays an important role in the early steps of CHC biosynthesis and impacts the abundance of methyl-branched alkanes (Chung et al. 2014; Wicker-Thomas et al. 2015). Additionally, *fas1* maps in close proximity to the conspicuous cluster of CHC biosynthesis gene orthologs (Fig. 5) that also co-localizes with QTL explaining variation of mostly methyl-branched alkanes between *N. vitripennis* and *N. giraulti* and between *N. longicornis* and *N. giraulti* (compare Fig. 3 and S1). This renders *fas1* a particularly promising candidate gene for further functional genetic studies investigating its impact on CHC profiles not only in *Nasonia*, but also in other Hymenoptera. Generally, establishing a functional link between the identified candidate genes and CHC variation will be instrumental in further elucidating the genomic and genetic architecture underlying species-specific CHC diversification in parasitoid wasps in particular and other hymenopteran species in general.

### Conclusions and Outlook

With our study on the genetic and genomic architecture of CHC variation between parasitoid wasps of the genus *Nasonia*, we uncovered several genomic regions where QTL explaining variation in similar CHC compounds and CHC biosynthesis-related candidate genes clustered. Especially one region on chromosome 1 close to the centromere is conspicuous in this regard, as it harbors many CHC biosynthesis-related candidate genes and QTL explaining variation of methyl-branched alkanes, the prevalent CHC compound class in the Hymenoptera in general and in *Nasonia* in particular. This underlines the considerable potential of *Nasonia* as a model system to further investigate the little-know genetic and genomic architecture of methyl-branched alkane biosynthesis and variation.

## ACKNOWLEDGEMENTS

Thanks to Andrea K. Judson, Nadine Brehm and Sylvia Geeritsma for their help in acquiring the chemical and QTL data, Joshua D. Gibson for his valuable input in conceptualizing the manuscript in its current form, and Valerio Vitali for helpful suggestions on the first draft of the manuscript.

## DATA AVAILABILITY STATEMENT

All data underlying the presented study will be made available at the dryad data repository

