## Supplementary material for "Genetic and genomic architecture of species-specific cuticular hydrocarbon variation in parasitoid wasps": Fig. S1

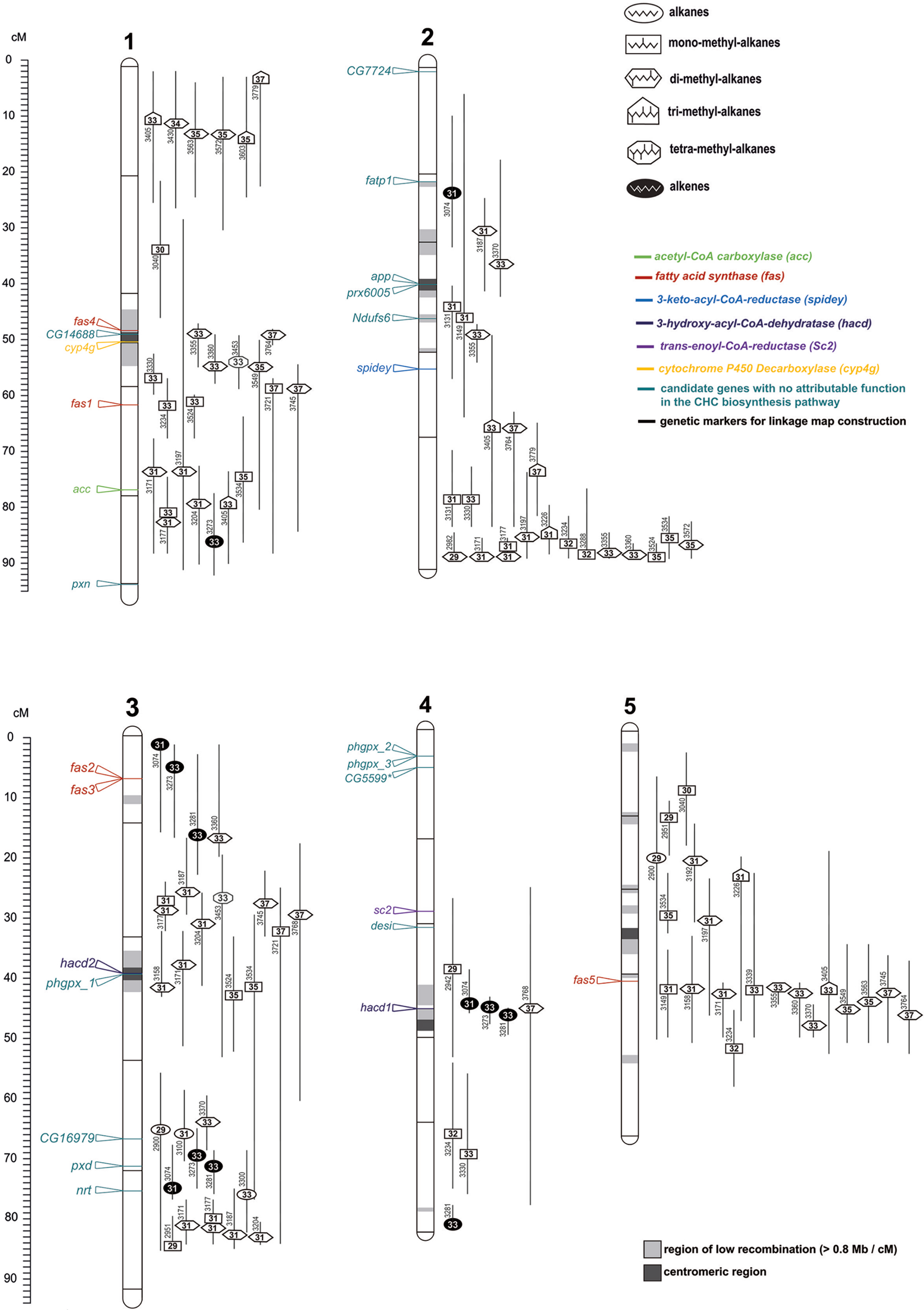

alkanes
mono-methyl-alkanes
di-methyl-alkanes
tri-methyl-alkanes
tetra-methyl-alkanes
alkenes
acetyl-CoA carboxylase (acc)
fatty acid synthase (fas)
3-keto-acyl-CoA-reductase (spidey)
3-hydroxy-acyl-CoA-dehydratase (hacd)
trans-enoyl-CoA-reductase (Sc2)
cytochrome P450 Decarboxylase (cyp4g)
candidate genes with no attributable function in the CHC biosynthesis pathway
genetic markers for linkage map construction
region of low recombination (> 0.8 Mb / cM)
centromeric region
cM
1
fas4
CG14688
cyp4g
fas1
acc
pxn
2
CG7724
fatp1
app
prx6005
Ndufs6
spidey
3
fas2
fas3
hacd2
phgpx_1
CG16979
pxd
nrt
4
phgpx_2
phgpx_3
CG5599*
sc2
desi
hacd1
5
fas5

**Figure S1**: Linkage map between *Nasonia vitripennis* (♂) and *Nasonia giraulti* (♀) depicting estimated positions of 102 predicted quantitative trait loci (QTL) in total for the variation of 42 individual cuticular hydrocarbons (CHCs) as well as mapped positions of CHC biosynthesis genes. CHCs are indicated by symbols corresponding to their respective compound class (n-alkanes, n-alkenes, mono-, di-, tri-, and tetra-methyl-alkanes) and their retention indices (RI). The five chromosomes are labeled 1–5. Labels of molecular markers and CHC biosynthesis gene orthologs as well as regions of low recombination (> 0.8 Mb / cM) and centromeric regions are indicated as in Fig. 3 and 4. Adapted from Niehuis et al., 2011.
