## Supplementary material for "Genetic and genomic architecture of species-specific cuticular hydrocarbon variation in parasitoid wasps": Tab. S1

**Tab. S1**: Length polymorphic molecular markers used to identify linked QTL positions associated with CHC variation between *Nasonia* hybrid males. Indicated are the respective Marker IDs based on the original scaffolds of the *Nasonia* genome assembly, the corresponding forward (F) and reverse (R) 5’→ 3’ PCR primer sequences, the respective amplicon lengths, and the positions of the markers in the *Nasonia* linkage map (centimorgan (cM) on the respective chromosome (c)).

| Marker ID | Primer sequences (5’→ 3’) | Amplicon length | Linkage map position |
| --- | --- | --- | --- |
| Scaf16_246267 | F: TCCGCACGGTCAGTCTTT  R: CGGCGAATTTCGTTCTTC | 119/109 | c1: 0 cM |
| Scaf16_3147035 | F: TTCCACTCGCTCTGTCCTAT  R: CGGAATACAGGACTCGATCCTA | 112/160 | c1: 19.7 cM |
| Scaf127_676512 | F: GGCTTCTCGCCATAATGAG  R: GCCGTATTGGCTCTCGTCTA | 178/216 | c1: 40.9 cM |
| Scaf1_1296440 | F: CGTGCACTTTCTCTCCCTTT  R: TGCACATTCGCGAAACAC | 242/258 | c1: 57.7 cM |
| Scaf1_3410194 | F: GCGCGTCGTCTTGTTTAA  R: TTACCCGGCCGATGTTAG | 196/204 | c1: 77.4 cM |
| Scaf7_288031 | F: GCTCGAGGAGGCCATATC  R: CCTCGATAGCTGGCAACC | 236/253 | c1: 93.4 cM |
| Scaf116_395861 | F: CGATGTTGCGACCGTCTATA  R: TCCGATCAAATCGAATTACTGTA | 255/234 | c2: 0.7 cM |
| Scaf3_1563474 | F: GCAGACACTGACACGTGATG  R: CCCTCGATCGCTGCACTA | 252/256 | c2: 19 cM |
| Scaf31_1706493 | F: GCACCGCTGCGATTAAAC  R: TGCTCTCGCTTCTCGAGTC | 271/299 | c2: 30.7 cM |
| Scaf19_1522711 | F: CCAACTTCTTATTCGTAAGGGAA  R: ACCATTCGCTGGCTGGTA | 193/205 | c2: 49.6 cM |
| Scaf21_835668 | F: AGGACGCAGCTAGGTGGC  R: CCTCGTCGATCAAGAGGC | 159/175 | c2: 64.2 cM |
| Scaf309_59696 | F: TGAAGAATGCGTATCAATCGTAC  R: ACCACCACGTCCTCCAAG | 139/175 | c2: 86.9 cM |
| Scaf18_70730 | F: CATAAATACATTTGGGTCTCCC  R: TGGAGTCCAGCTAGGATTCTAA | 177/191 | c3: 0 cM |
| Scaf42_535846 | F: GGAAGGAGCGAATCCTCTAC  R: AGTGCGTCTCGACGCTAG | 234/305 | c3: 13.9 cM |
| Scaf6_1979404 | F: GCGAGCAGTGCGAGATTA  R: CGGTTGTACTTTGGCGATAAA | 242/250 | c3: 32.1 cM |
| Scaf22_2413317 | F: CACGAAACTACATCGCAATCA  R: CGTGTATAGCTGCTCTTGTTGAA | 294/333 | c3: 51.8 cM |
| Scaf17_2027794 | F: CGCTCTCTGCGTGTGTCT  R: AGAGCGATAGCGTCGCTT | 228/262 | c3: 69.4 cM |
| Scaf28_1519494 | F: AATGGCATTATGCGAATGA  R: CTGCTCTCTGCATGAATCTTT | 202/219 | c3: 88.3 cM |
| Scaf4_4991034 | F: TCGCTTAGATAATTGCCAGAC  R: ACAGATATACTCTCGTGCAGGAG | 190/161 | c4: 0 cM |
| Scaf4_2173128 | F: GCCTGCCGTACAATCAAA  R: GAAACGCGACGCTGTTAG | 225/235 | c4: 19.7 cM |
| Scaf4_148046 | F: CGCTACTCTCGTCAACCTGTAA  R: GTTCGCTCGTCGATGATAA | 290/314 | c4: 35 cM |
| Scaf40_1810242 | F: GTCCGTGTGTACTGCGAAG  R: ACCTCGGAAACGGCTAGA | 172/180 | c4: 55.5 cM |
| Scaf9_3686475 | F: TCCTTTCTCGCAAATTGTTAA  R: GCGCCGCTACAGTTAGAC | 267/245 | c4: 70.8 cM |
| Scaf9_46733 | F: CGGATATCTACGGAAATAGCATT  R: AACACACTCGCTCGCTTT | 222/250 | c4: 90.6 cM |
| Scaf14_13473 | F: CGGCATACCTATAGCGCAGA  R: CCATCCATTCGGAATACAATCT | 296/270 | c5: 0 cM |
| Scaf38_564492 | F: GGATTCGCTCCGCAATTA  R: CATTTCCGCGTCCCTCTT | 243/259 | c5: 15.3 cM |
| Scaf1_5852676 | F: GTGACGCTTCGCAGCTGT  R: GGATATTCGCGGCGAGTG | 277/301 | c5: 28.5 cM |
| Scaf176_257755 | F: GATGGACCGATTCGCACA  R: CGAGAGGTATACACCTGGTGGT | 99/77 | c5: 43.8 cM |
| Scaf2_252533 | F: AAATATTGGCGCGGCAAC  R: CCAACAATGAGTGTATCCTAGGC | 225/255 | c5: 73 cM |
