## Supplementary material for "Genetic and genomic architecture of species-specific cuticular hydrocarbon variation in parasitoid wasps": Tab. S2

**Tab. S2**: Candidate genes from *Drosophila* *melanogaster* with an unambiguously demonstrated effect on CHC profile compositions, selected to screen for orthologs in the *Nasonia vitripennis* reference genome. Indicated are gene annotations and IDs obtained from NCBI, gene IDs and (predicted) molecular function of the respective gene product from FlyBase (Version FB2019_02, Gramates et al. 2017), description of the concrete effects on *D. melanogaster* CHC profiles, references, and the number of orthologs obtained via three different ortholog screening methods (Reciprocal BLAST search: RB, OrthoFinder: OF, WaspAtlas: WA) as well as an indicator whether the overall evidence of orthology was strong enough for further consideration for each gene. (+) indicates one unambiguously identified ortholog by all three methods, (-) indicates no clearly detectable orthologs in the *N. vitripennis* reference genome, (ǂ) indicates discrepancies between the three orthology inference methods or unclear gene phylogenies, in which case we did not consider those candidate genes further for our analysis. In cases where further investigation led to acceptance of single candidate genes for further analysis, the most reliable method of orthology inference is indicated. Orthogroups from the OrthoFinder (OF) orthology inference method that contained multiple *D. melanogaster* genes are marked with an asterisk.

| NCBI annotation | NCBI gene ID | FlyBase Gene ID | (Predicted) molecular function in FlyBase | Effect on CHC profiles | Reference | Ortholog hits per method | | | Orthology evidence |
| --- | --- | --- | --- | --- | --- | --- | --- | --- | --- |
|  |  |  |  |  |  | RB | OF | WA |  |
| *acc* | 35761 | 33246 | acetyl-CoA carboxylase activity; ATP binding; metal ion binding | almost full depletion of all CHCs | Wicker-Thomas et al, 2015 | 1 | 1 | 1 | + |
| *app* | 39399 | 260941 | protein-cysteine S-palmitoyltransferase activity | unspecific upregulation of CHCs, female-specific downregulation of 1 n-alkene | Dembeck et al, 2015 | 1 | 1 | 1 | + |
| *CG10097* | 3771756 | 38033 | fatty-acyl-CoA reductase (alcohol-forming) activity | unspecific up- and downregulation of CHCs | Dembeck et al, 2015 | - | - | - | - |
| *CG13091* | 34188 | 32055 | fatty-acyl-CoA reductase (alcohol-forming) activity | unspecific up- and downregulation of CHCs | Dembeck et al, 2015 | - | - | - | - |
| *CG14688* | 41275 | 37819 | phytanoyl-CoA dioxygenase activity | unspecific upregulation of CHCs in females, unspeci-fic up- and downregula-tion of CHCs in males (no downregulation of n-alkanes) | Dembeck et al, 2015 | 1 | 1 | 1 | + |
| *CG16979* | 39687 | 36512 | thiolester hydrolase activity; UFM1 hydrolase activity | unspecific upregulation of CHCs in males only | Dembeck et al, 2015 | 1 | 1 | 1 | + |
| *CG18609* | 37158 | 34382 | fatty acid elongase activity | unspecific up- and downregulation of CHCs | Dembeck et al, 2015 | - | - | 1 | ǂ |
| *CG30008* | 246388 | 50008 | fatty acid elongase activity | unspecific up- and downregulation of CHCs, much stronger in males | Dembeck et al, 2015 | - | - | 4 | ǂ |
| *CG5599* | 32441 | 30612 | acetyltransferase activity; dihydrolipoamide branched chain acyltransferase activity; lipoic acid binding | upregulation of all CHCs | Dembeck et al, 2015 | 1 | 1 | 1 | + |
| *CG7724* | 39918 | 36698 | 3-beta-hydroxy-delta5-steroid dehydrogenase activity; oxidoreductase activity; acting on the CH-OH group of donors, NAD or NADP as acceptor; steroid delta- isomerase activity | unspecific up- and downregulation of CHCs | Dembeck et al, 2015 | 1 | 1 | 1 | + |
| *CG8680* | 33744 | 31684 | - | Upregulation of all CHCs | Dembeck et al, 2015 | 1 | 1 | 1 | + |
| *CG8814* | 33492 | 31478 | fatty-acyl-CoA binding | unspecific upregulation of CHCs | Dembeck et al, 2015 | - | - | - | - |
| *CG9458* | 41214 | 37765 | fatty acid elongase activity | unspecific downregulation of CHCs in females (upregulation of 1 mb-alkane), unspecific upregulation of CHCs in males | Dembeck et al, 2015 | - | - | 1 | - |
| *CG9801* | 41044 | 37623 | catalytic activity | unspecific downregulation of CHCs and upregulation of 1 mb-alkane in females, unspecific upregulation of CHCs in males | Dembeck et al, 2015 | - | - | - | - |
| *Cyp49a1* | 36105 | 33524 | heme binding; iron ion binding; oxidoreductase activity, acting on paired donors, with incorporation or reduction of molecular oxygen | unspecific downregulation of CHCs in females, unspecific upregulation of CHCs in males | Dembeck et al, 2015 | 1 | * | 1 | ǂ |
| *Cyp4g1* | 30986 | 10019 | aldehyde decarbonylase activity | Strong downregulation of all CHC CHCs after 4 days, retention of longer chained CHC after 1 day | Qiu et al, 2012 | - | - | 1 | WA |
| *Cyp4s3* | 32444 | 30615 | heme binding; iron ion binding; oxidoreductase activity, acting on paired donors, with incorporation or reduction of molecular oxygen | unspecific upregulation of CHCs in females, unspeci-fic up- and down-regulation of CHCs in males | Dembeck et al, 2015 | - | - | - | - |
| *Cyp9f2* | 41520 | 38037 | heme binding; iron ion binding; oxidoreductase activity, acting on paired donors, with incorporation or reduction of molecular oxygen | unspecific up- and downregulation of CHCs in females, unspecific upregulation of CHCs in males | Dembeck et al, 2015 | 16 | * | 15 | ǂ |
| *Desat1* | 117369 | 86687 | iron ion binding; stearoyl-CoA 9-desaturase activity | Downregulation of both n-n-alkenes and alkadienes in males and females | Wicker-Thomas et al, 1997; Dallerac et al, 2000 | 4 | * | 1 | ǂ |
| *Desat2* | 41536 | 43043 | iron ion binding; stearoyl-CoA 9-desaturase activity | Conversion of 1 specific diene compound to its positional isomer | Coyne et al, 1999; Dallerac et al, 2000 | 1 | * | 1 | ǂ |
| *DesatF* | 44006 | 29172 | iron ion binding; stearoyl-CoA 9-desaturase activity | Knockdown effect only in females, upregulation of n-alkenes, downregulation of alkadienes | Dembeck et al, 2015 | - | - | - | - |
| *Desi* | 41292 | 37832 | PDZ domain binding; SH2 domain binding | unspecific upregulation of CHCs, only in males | Chertemps et al, 2007 | 1 | 1 | 1 | + |
| *eloF* | 41211 | 37762 | fatty acid elongase activity | Only females: Large increase of shorter- chain CHCs, decrease of longer-chain CHCs | Chertemps et al, 2006 | - | - | 1 | ǂ |
| *FASN1* | 33524 | 283427 | fatty acid synthase activity | fat body-expressed | Wicker-Thomas et al, 2015 | 2 | (5) | - | OF |
| *FASN2/mFAS* | 117361 | 42627 | fatty acid synthase activity | oenocyte-specific, putative microsomal FAS, effect on males only | Chung et al, 2014; Wicker-Thomas et al, 2015 | - | (5) | 1 | OF |
| *FASN3* | 3355111 | 40001 | fatty acid synthase activity; hydrolase activity, acting on ester bonds; oxidoreductase activity; phosphopantetheine binding | oenocyte-specific, putative cytosolic FAS, effect on males only | Wicker-Thomas et al, 2015 | 7 | (5) | 2 | OF |
| *Fatp1* | 2.6E+07 | 267828 | long-chain fatty acid transporter activity; long-chain fatty acid-CoA ligase activity; very long- chain fatty acid-CoA ligase activity | unspecific downregula-tion of CHCs, alkadienes only detected and downregulated in females | Wicker-Thomas et al, 2015 | 1 | * | - | RB |
| *Hacd1* | 34614 | 32394 | 3-hydroxyacyl-CoA dehydratase activity;enzyme binding | unspecific downregula-tion of CHCs, alkadienes only detected and downregulated in females | Wicker-Thomas et al, 2015 | 1 | 1 | 1 | + |
| *Hacd2* | 34762 | 32524 | 3-hydroxyacyl-CoA dehydratase activity; enzyme binding | unspecific downregulation of CHCs, alkadienes only detected and downregulated in females, mb-alkanes only in males | Wicker-Thomas et al, 2015 | 1 | 1 | 1 | + |
| *Irc* | 42049 | 38465 | catalase activity | unspecific downregulation of CHCs in females (except for one diene), unspecific upregulation of CHCs in males | Dembeck et al, 2015 | - | - | - | - |
| *Lip2* | 43980 | 24740 | lipase activity | upregulation of mostly n-alkanes and methyl-branched alkanes, downregulation of only n-alkanes and n-alkenes in males | Dembeck et al, 2015 | - | - | - | - |
| *Nrt* | 39873 | 4108 | carboxylic ester hydrolase activity | unspecific up- and downregulation of CHCs in females, unspecific upregulation of CHCs and downregulation of only n-alkanes in males | Dembeck et al, 2015 | 1 | 1 | 1 | + |
| *PHGPx* | 38413 | 35438 | peroxidase activity | unspecific up- and downregulation of CHCs in females, unspecific upregulation of CHCs in males | Dembeck et al, 2015 | 3 | 1 | 3 | RB, WA |
| *Prx6005* | 33493 | 31479 | peroxidase activity | unspecific up- and downregulation of CHCs | Dembeck et al, 2015 | 1 | 1 | 1 | + |
| *Pxd* | 2768671 | 4577 | heme binding; peroxidase activity | unspecific upregulation of CHCs in females, unspecific up- and downregulation of CHCs in males | Dembeck et al, 2015 | 3 | 1 | 1 | OF, WA |
| *Pxn* | 38326 | 11828 | heme binding; peroxidase activity | unspecific up- and downregulation of CHCs | Dembeck et al, 2015 | 1 | 1 | 1 | + |
| *Sc2/TER* | 38457 | 35471 | oxidoreductase activity; oxidoreductase activity, acting on the CH-CH group of donors | unspecific, strong downregulation of CHCs, alkadienes only detected and downregulated in females | Wicker-Thomas et al, 2015 | 1 | 1 | 1 | + |
| *spidey/KAR* | 31703 | 29975 | - | almost full depletion of all CHCs, alkadienes only detected and depleted in females | Wicker-Thomas et al, 2015 | 1 | 1 | 1 | + |
