## Supplementary material for "Genetic and genomic architecture of species-specific cuticular hydrocarbon variation in parasitoid wasps": Tab. S3

**Tab. S3**: Male *Nasonia* cuticular hydrocarbons (CHCs) identifications and the corresponding QTL governing their variation between male hybrids obtained from crosses between *N. vitripennis* and *N. girauli* (Niehuis et al., 2011). Indicated are the retention indices (RI), compound identifications or possible configurations in case of ambiguities, chromosome number (Chr.), QTL position (pos.), confidence interval for the respective QTL position, ratios of the CHC compounds in parental males of the two species (percentages with standard deviations) and significance assessments comparing the two ratios (Benjamini Hochberg-corrected Mann-Whitney U tests).

| RI | Compound identification / possible configurations | Chr. | QTL pos. | Confidence interval | NV ♂ ratio | NG ♂ ratio | Significance |
| --- | --- | --- | --- | --- | --- | --- | --- |
| 2900 | C29 | 3 | 70 | 59.7 - 92.1 | 0.98 ± 0.5 | 1.23 ± 0.36 | n.s. |
|  |  | 5 | 23 | 8 - 56 |  |  |  |
| 2942 | 7-MeC29 | 4 | 41 | 28 - 57 | 0.6 ± 0.31 | 0.13 ± 0.17 | 0.000462 |
| 2951 | 5-MeC29 | 3 | 92 | 86 - 92.1 | 0.37 ± 0.16 | 0.16 ± 0.13 | 0.011683 |
|  |  | 5 | 16 | 13–23 |  |  |  |
| 2982 | 15,17-DiMeC29 | 2 | 89.8 | 85 - | 0.15 ± 0.07 | 0.01 ± 0.03 | 0.000184 |
| 3040 | 7-MeC30 | 1 | 33 | 20 - 45 | 0.33 ± 0.11 | 0.14 ± 0.1 | 0.001215 |
|  |  | 5 | 11 | 4–21 |  |  |  |
| 3049 | 5-MeC30 | 5 | 16 | 13 - 23 | 0.05 ± 0.03 | 0.01 ± 0.02 | n.s. |
| 3074 | 9-C31ene | 2 | 23 | 9 - 32 | 0.98 ± 0.73 | 0.08 ± 0.1 | n.s. |
|  |  | 3 | 16 | 0 - |  |  |  |
|  |  | 3 | 81 | 73 - 83 |  |  |  |
|  |  | 4 | 47 | 41 - 49 |  |  |  |
| 3100 | C31 | 3 | 71 | 63 - 76 | 6.41 ± 1.54 | 13 ± 2.96 | < 0.0001 |
| 3131 | 9-; 11-; 13-; 15-MeC31 | 2 | 44.2 | 40 - 57 |  |  |  |
|  |  | 2 | 79 | 70 - 84 |  |  |  |
| 3141 | 7-MeC31 | 5 | 11 | 4 - 21 | 11.43 ± 1.11 | 10.78 ± 1.79 | n.s. |
| 3149 | 5-MeC31 | 2 | 46 | 5 - 64 | 4.42 ± 0.65 | 5.68 ± 0.81 | 0.00204 |
|  |  | 5 | 47 | 40 - 56 |  |  |  |
| 3158 | 11,15-; 11,17-; 11,21-DiMeC31 | 3 | 44 | 34 - 46 | 0.46 ± 0.13 | 0.24 ± 0.05 | 0.00204 |
|  |  | 5 | 47.6 | 37.7 - 57 |  |  |  |
| 3171 | 7,11-DiMeC31 | 1 | 73 | 67 - 88.1 | 3.73 ± 0.35 | 4.72 ± 0.74 | 0.000306 |
|  |  | 2 | 89.8 | 86 - |  |  |  |
|  |  | 3 | 40 | 34 - 55 |  |  |  |
|  |  | 3 | 88 | 83 - 91.2 |  |  |  |
|  |  | 5 | 48 | 46 - 56 |  |  |  |
| 3177 | 3-MeC31 (+7,23-DiMeC31) | 1 | 82 | 74 - 88.1 | 2.55 ± 0.55 | 1.44 ± 0.16 | < 0.0001 |
|  |  | 2 | 89.8 | 86 - |  |  |  |
|  |  | 3 | 30 | 25 - 34 |  |  |  |
|  |  | 3 | 88 | 83 - 91.2 |  |  |  |
| 3187 | 7,25-DiMeC31 | 2 | 30 | 24 - 41 | 0.61 ± 0.18 | 0.19 ± 0.05 | < 0.0001 |
|  |  | 3 | 27 | 17 - 31 |  |  |  |
|  |  | 3 | 89 | 81 - 92.1 |  |  |  |
| 3192 | 3,25-DiMeC31 | 5 | 24 | 17 - 35 | 0.34 ± 0.09 | 0.48 ± 0.08 | 0.00174 |
| 3197 | 3,15-DiMeC31 | 1 | 73 | 27 - 91 | 0.22 ± 0.05 | 0.43 ± 0.12 | < 0.0001 |
|  |  | 2 | 86 | 74 - 89.8 |  |  |  |
|  |  | 5 | 35 | 27 - 52 |  |  |  |
| 3204 | 3,7-DiMeC31 | 1 | 79 | 72 - 90 | 0.81 ± 0.17 | 0.42 ± 0.06 | < 0.0001 |
|  |  | 3 | 33 | 27 - 44 |  |  |  |
|  |  | 3 | 90 | 87 - 91.2 |  |  |  |
| 3226 | 3,7,9-; 3,7,11-; 3,7,15-TriMeC31 | 2 | 85 | 81 - 89 | 0.16 ± 0.08 | 0.27 ± 0.09 | 0.025044 |
|  |  | 5 | 27 | 23 - 53 |  |  |  |
| 3234 | 6-MeC32 | 1 | 61 | 56 - 67 | 0.39 ± 0.05 | 0.51 ± 0.12 | n.s. |
|  |  | 2 | 87 | 82 - 89.8 |  |  |  |
|  |  | 4 | 71 | 58 - 81 |  |  |  |
|  |  | 5 | 58 | 51 - 65 |  |  |  |
| 3273 | 9-C33ene | 1 | 86 | 77 - 92 | 1.45 ± 0.27 | 0.16 ± 0.13 | < 0.0001 |
|  |  | 3 | 4 | 0 - 17 |  |  |  |
|  |  | 3 | 75 | 70 - 81 |  |  |  |
|  |  | 4 | 48 | 46 - 51 |  |  |  |
| 3281 | 7-C33ene | 3 | 17 | 2 - 24 | 0.96 ± 0.2 | 0.06 ± 0.1 | < 0.0001 |
|  |  | 3 | 77 | 74 - 82 |  |  |  |
|  |  | 4 | 49 | 48 - 53 |  |  |  |
|  |  | 4 | 87.6 | 84 - |  |  |  |
| 3288 | 3-MeC32 | 2 | 89.8 | 77 - | 0.12 ± 0.04 | 0.18 ± 0.09 |  |
| 3300 | C33 | 3 | 82 | 74 - 89 | 0.99 ± 0.25 | 1.38 ± 0.41 | n.s. |
| 3330 | 9-; 11-; 13-; 15-MeC33 | 1 | 56 | 51.6 - 59 | 3.97 ± 1.26 | 8.02 ± 0.92 | < 0.0001 |
|  |  | 2 | 79 | 73 - 84 |  |  |  |
|  |  | 4 | 74.5 | 60 - 82 |  |  |  |
| 3339 | 7-MeC33 | 5 | 47.6 | 26 - 56 | 2.93 ± 0.68 | 3.54 ± 0.8 | n.s. |
| 3355 | 11,15-; 11,23- 11,25-DiMeC33 | 1 | 48 | 46 - 54 | 0.66 ± 0.42 | 2.29 ± 0.64 | < 0.0001 |
|  |  | 2 | 49 | 47 - 54 |  |  |  |
|  |  | 2 | 89 | 85 - 89.8 |  |  |  |
|  |  | 5 | 50 | 47 - |  |  |  |
| 3360 | 7,19-; 7,21-DiMeC33 | 1 | 54 | 48 - 57.1 | 1.68 ± 0.56 | 1.24 ± 0.21 | n.s. |
|  |  | 2 | 89.8 | 87 - |  |  |  |
|  |  | 3 | 17 | 0 - 20.4 |  |  |  |
|  |  | 5 | 48 | 46 - 56 |  |  |  |
| 3370 | 7,23-DiMeC33 | 2 | 36 | 17 - 42 | 7.71 ± 1.21 | 3.24 ± 0.38 | < 0.0001 |
|  |  | 3 | 69 | 64 - 74 |  |  |  |
|  |  | 5 | 54 | 50 - 56 |  |  |  |
| 3405 | 5,9,11-; 5,9,15-TriMeC33 | 1 | 9 | 0 - 24 | 2.04 ± 0.68 | 0 | < 0.0001 |
|  |  | 1 | 79 | 73 - 89.9 |  |  |  |
|  |  | 2 | 66 | 49 - 84 |  |  |  |
|  |  | 5 | 47 | 22 - 59 |  |  |  |
| 3430 | 8,10-; 8,12-; 8,14-; 8,16-; 8,18-DiMeC34 | 1 | 10 | 0 - 25 | 0.92 ± 0.28 | 0.84 ± 0.21 | n.s. |
| 3453 | 3,7,11,15-TetraMeC33 | 1 | 53.4 | 48 - 58 | 1.57 ± 0.35 | 4 ± 0.71 | < 0.0001 |
|  |  | 3 | 28 | 20 - 57 |  |  |  |
| 3524 | 13-; 15-;17-MeC35 | 1 | 60 | 59 - 67 | 2.58 ± 0.44 | 3.34 ± 0.38 | 0.000636 |
|  |  | 2 | 89.8 | 87 - |  |  |  |
|  |  | 3 | 46 | 35 - 56 |  |  |  |
| 3534 | 7-MeC35 | 1 | 74 | 63 - 86 | 0.77 ± 0.18 | 0.77 ± 0.12 | n.s. |
|  |  | 2 | 86 | 82 - 89.8 |  |  |  |
|  |  | 3 | 44.3 | 31 - 83 |  |  |  |
|  |  | 5 | 34 | 26 - 37 |  |  |  |
| 3549 | 11,17-; 11,19-; 11,21-; 11,23-; 13,17-; 13,19-; 13,21-; 13,17-; 13,19-; 13,21-;13,23-; 15,17-; 15,19-; 15,21-; 15,23-DiMeC35 | 1 | 54 | 49 - 80 | 3.08 ± 0.92 | 6.55 ± 0.86 | < 0.0001 |
|  |  | 5 | 51 | 39 - 57 |  |  |  |
| 3563 | 7,19-; 7,21-; 7,23-DiMeC35 | 1 | 12 | 2 - 23 | 7.07 ± 1.47 | 2.15 ± 0.28 | < 0.0001 |
|  |  | 5 | 46 | 39 - 57 |  |  |  |
| 3572 | 5,15-; 5,17-; 5,19-; 5,21-; 5,23-DiMeC35 | 1 | 12 | 1 - 29 | 5.21 ± 0.73 | 3.54 ± 0.53 | < 0.0001 |
|  |  | 2 | 88 | 83 - 89.8 |  |  |  |
| 3603 | 5,9,13-; 5,9,15-; 5,9,17-; 5,9,19-; 5,9,21-; 5,11,13-; 5,11,15-; 5,11,17-; 5,11,19-; 5,11,21-TriMeC35 | 1 | 13 | 1 - 23 | 1.65 ± 0.4 | 1.05 ± 0.33 | 0.00324 |
| 3721 | 13-; 15-; 17-; 19-MeC37 | 1 | 58 | 56 - 88 | 0.64 ± 0.23 | 0.9 ± 0.27 |  |
|  |  | 3 | 34 | 26 - 91 |  |  |  |
| 3745 | 11,19-; 11,21-; 11,23-; 11,25-; 13,19-; 13,21-; 13,23-; 15,19-; 15,21-; 15,23-DiMeC37 | 1 | 58 | 53.4 - 84 | 1.38 ± 0.46 | 2.13 ± 0.38 | 0.010146 |
|  |  | 3 | 29 | 23 - 35 |  |  |  |
|  |  | 5 | 48 | 41 - 57 |  |  |  |
| 3764 | 7,19-; 7,21-; 7,23-DiMeC37 | 1 | 48 | 47 - 50 | 2.39 ± 0.69 | 0.61 ± 0.16 | < 0.0001 |
|  |  | 2 | 66 | 63 - 84 |  |  |  |
|  |  | 5 | 52 | 42 - 59 |  |  |  |
| 3768 | 5,15-; 5,17-; 5,23-DiMeC37 | 3 | 31 | 18 - 65 | 1.42 ± 0.35 | 1.31 ± 0.28 | n.s. |
|  |  | 4 | 48.2 | 26 - 84 |  |  |  |
| 3779 | 7,15,17-; 7,15,19-; 7,15,23-; 7,19,17-; 7,19,19-; 7,19,23-TriMeC37 | 1 | 1 | 0 - 21 | 0.53 ± 0.36 | 0.03 ± 0.09 | 0.00016 |
|  |  | 2 | 74 | 66 - 82 |  |  |  |
