## Supplementary material for "Genetic and genomic architecture of species-specific cuticular hydrocarbon variation in parasitoid wasps": Tab. S4

**Tab. S4**: Average CHC quantities (percentages and standard deviations) per individual compound class (n-alkanes, n-alkenes, mono-methyl-alkanes, di-methyl-alkanes, tri-methyl-alkanes and tetra-methyl-alkanes) detected in males of the three *Nasonia* species *N. giraulti* (NG), *N. longicornis* (NL) and *N. vitripennis* (NV). Quantities and standard deviations are summed up for methyl-branched alkanes in total. Sample sizes: NG = 12, NL = 11, NV = 32.

|  | NG ♂ ratio | NL ♂ ratio | NV ♂ ratio |
| --- | --- | --- | --- |
| n-alkanes | 16.15 ± 3.5 | 8.98 ± 2.75 | 8.73 ± 2.15 |
| n-alkenes | 0.29 ± 0.3 | 0.52 ± 0.24 | 3.41 ± 0.91 |
| mono-methyl alkanes | 38.84 ± 3.44 | 50.25 ± 3.97 | 34.77 ± 3.3 |
| di-methyl alkanes | 37.67 ± 4.34 | 37.02 ± 5.22 | 45.41 ± 4.01 |
| tri-methyl alkanes | 2.36 ± 0.57 | 1.81 ± 0.57 | 5.77 ± 1.63 |
| tetra-methyl alkanes | 4.69 ± 0.87 | 1.42 ± 0.71 | 1.9 ± 0.42 |
| methyl alkanes total | 83.56 ± 3.45 | 90.5 ± 2.63 | 87.85 ± 2.74 |
