## Supplementary material for "Genetic and genomic architecture of species-specific cuticular hydrocarbon variation in parasitoid wasps": Tab. S5

**Tab. S5**: Average CHC quantities (percentages and standard deviations) per individual compound class (n-alkanes, n-alkenes, alkadienes, mono-methyl-alkanes and unknown) summarized for males and females of the lines in the *Drosophila melanogaster* Reference Panel (DGRP), obtained from Dembeck et al., 2015.

|  | DM ♂ ratio | DM ♀ ratio |
| --- | --- | --- |
| n-alkanes | 24.91 ± 6.18 | 25.69 ± 5.84 |
| n-alkenes | 58.51 ± 7.5 | 8.26 ± 4.03 |
| dienes | 0.67 ± 0.34 | 40.55 ± 7.57 |
| mono-methyl alkanes | 16.15 ± 4.37 | 24.13 ± 5.83 |
| unknown | 0.39 ± 0.14 | 0.49 ± 0.21 |
